# Cell surface receptors TREM2, CD14 and integrin α_M_β_2_ drive sinking engulfment in phosphatidylserine-mediated phagocytosis

**DOI:** 10.1101/2022.07.30.502145

**Authors:** Daan Vorselen, Roarke A. Kamber, Ramon Lorenzo D. Labitigan, Aaron P. van Loon, Eric Peterman, Melissa K. Delgado, Sijie Lin, Jeffrey P. Rasmussen, Michael C. Bassik, Julie A. Theriot

**Affiliations:** Department of Biology, University of Washington, Seattle, WA, 98105, USA; Howard Hughes Medical Institute, University of Washington, Seattle, WA, 98105, USA; Department of Genetics, Stanford University School of Medicine, Stanford, CA, 94305, USA; Department of Biochemistry, Stanford University School of Medicine, Stanford, CA, 94305, USA

**Keywords:** Macrophages, phagocytosis, phosphatidylserine, CRISPR screen, TREM2

## Abstract

Macrophages phagocytose and thereby eliminate a wide array of extracellular threats, ranging from antibody-coated bacteria to apoptotic cells. Precision modulation of phagocytosis has emerged as a therapeutic strategy across a range of diseases, but is limited by our incomplete understanding of how macrophages recognize, engulf, and respond to different phagocytic targets. Here, we undertook a systematic investigation of the morphological, biophysical and regulatory differences between two major types of phagocytosis: an immunostimulatory form of phagocytosis triggered by antibody-coated targets and an immunosuppressive form triggered by phosphatidylserine (PS)-coated targets. We confirmed classic observations that antibody-mediated phagocytosis involves the extension of thin actin-rich protrusions around the target, but find that PS-mediated phagocytosis involves an unexpected combination of filopodial probing, piecemeal phagocytosis and a distinct ‘sinking’ mechanism of uptake. Using a genome-wide screening approach, we identified genes specifically required for each form of phagocytosis, including actin regulators, cell surface receptors and intracellular signaling molecules. Three cell surface receptors - TREM2, CD14 and integrin α_M_β_2_ - were revealed as essential for PS-mediated uptake. Strikingly, each receptor exhibited a distinct pattern of localization at the plasma membrane and contributed uniquely to the organization of the PS-dependent phagocytic cup. Overall, this work reveals divergent genetic requirements for the morphologically and mechanically distinct forms of PS-mediated and antibody-mediated phagocytosis, thereby informing therapeutic strategies for substrate-specific phagocytosis modulation.

## Introduction

Macrophages detect and engulf via phagocytosis a wide range of large (>0.5 µm in diameter) substrates, ranging from dead cells and cellular debris to bacteria and antibody-coated cancer cells. Phagocytosis is critical for organismal health not only by clearing disease-causing cells, pathogens and extracellular material, but by maintaining tissue homeostasis via appropriate tissue-level responses based on sensing the nature of the engulfed material. Whereas phagocytosis of targets not associated with infection (e.g. apoptotic cells) has been shown to be “immunologically silent”, phagocytosis of infection-associated material (e.g. antibody-coated bacteria) stimulates release of inflammatory cytokines and presentation of antigens derived from the engulfed material (Yin and Heit, 2021), (Aderem and Underhill, 1999; Fadok et al., 1998). How different opsonins drive these different macrophage responses remains largely unclear. In disease states such as cancer, atherosclerosis, and neurodegeneration, uptake of disease-associated substrates (such as an antibody-coated cancer cell within the tumor mass, or dead cells in atherosclerotic plaques) has been variably reported to drive both subsequent immune tolerance and an inflammatory response (Bäck et al., 2019; Boada-Romero et al., 2020; Heckmann et al., 2017; Su et al., 2018). Understanding the molecular basis for these disparate responses will require an integrative understanding of the signaling pathways and ultrastructural processes linking recognition, uptake, and response to different phagocytic substrates. Two key barriers to our understanding are: 1) our incomplete knowledge of the receptors that mediate recognition of immunogenic vs non-immunogenic targets and 2) our lack of knowledge of the possible morphological and mechanical differences between immunogenic and non-immunogenic modes of phagocytosis.

Here we undertook a systematic study of both the genetic and morphological distinctions between phagocytosis of targets opsonized with a non-immunogenic ligand (phosphatidylserine) and an immunogenic ligand (IgG). Surprisingly, we find that uptake of apoptotic cells and phosphatidylserine-coated beads involves a distinct mode of engulfment from canonical phagocytosis of antibody-coated targets, which involves protrusion of pseudopods from the macrophage to engulf the target. By contrast, apoptotic cells are phagocytosed via a combination of trogocytosis (piecemeal uptake) and “sinking” phagocytosis, in which the apoptotic cell appears sunken into the cytoplasm before engulfment has been completed. Comparative genome-wide genetic screens for regulators of PS-driven sinking phagocytosis and IgG-driven reaching phagocytosis revealed a striking convergence in the requirement for a core set of actin regulators for both forms of phagocytosis. At the same time, these screens also revealed exquisite specificity for cell surface receptors and intracellular signaling molecules for each form of phagocytosis. Interestingly, three previously proposed phagocytic receptors, TREM2, CD14, and integrin α_M_β_2_ (Mac-1) (Deczkowska et al., 2020; Freeman and Grinstein, 2014), were uncovered as playing non-redundant roles in uptake of PS-coated particles. Using super-resolution microscopy and quantitative image analysis, we found that TREM2 localizes to the rim of the phagocytic cup, and is critical for inducing formation of a sinking phagocytic cup.

## Results

### Apoptotic Cell Uptake Involves F-actin-rich Protrusions, Sinking, and Partial Phagocytosis

To discover the dynamics of phagocytosis of apoptotic cells (efferocytosis), we induced apoptosis in B-cell lymphoma derived A20 cells using 20 h of treatment with 5 μM staurosporine, which led to a strong (> 3-fold) increase of apoptotic cell marker phosphatidylserine (PS) on the cell surface (Fig. S1A). This treatment also led to increased uptake of A20s when they were fed to J774 macrophages transduced with LifeAct-EGFP (Fig. S1B). Phagocytosis of apoptotic cells often occurred through partial uptake, reminiscent of trogocytosis, or “nibbling” (Zhao et al., 2022). Notably, trogocytosis was observed together with macrophage cell surface protrusions strongly enriched in F-actin protruding deep (∼10 μm) into the apoptotic cells (Fig. 1A, white arrows). Whole-cell phagocytosis also occurred, although rarely (∼6% of phagocytic cells; Fig. S1D). Our results are consistent with recent observations of partial apoptotic cell uptake by epithelial cells in the zebrafish embryo, and suggest that trogocytosis may be more general in efferocytosis (Hoijman et al., 2021).

**Figure 1:**
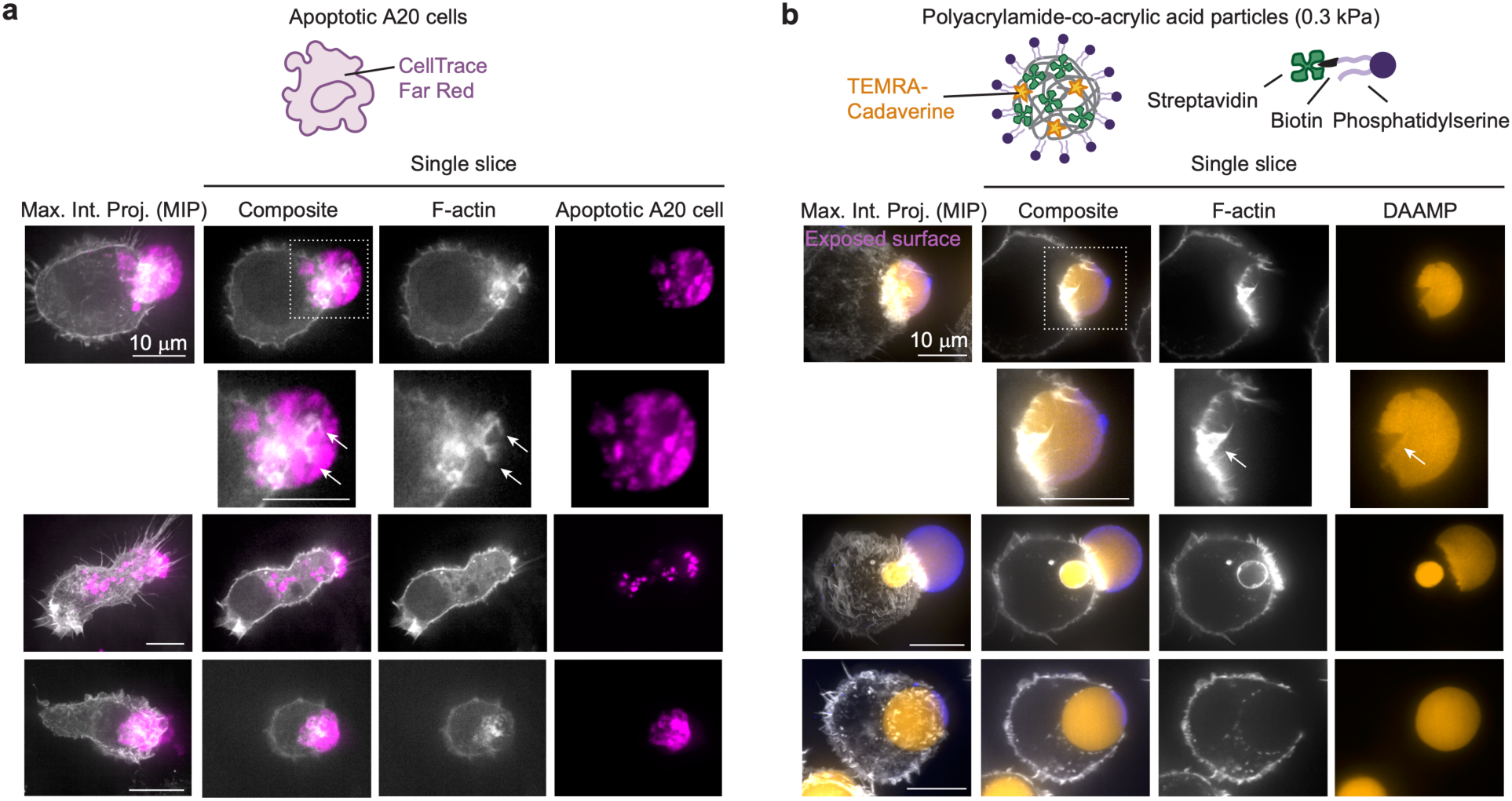
PS-mediated uptake of apoptotic cells and particles involves protrusive F-actin structures, trogocytosis and sinking phagocytosis. **a,** Super-resolution iSIM images of fixed J774 macrophages, which were transfected with LifeAct-EGFP, interacting with apoptotic A20 cells, which were treated with 5 μM staurosporine for 20 h and stained with CellTrace Far Red. Images were acquired 45 or 90 minutes after bringing the cells in contact. Left column: composite maximum intensity projections (MIP) of z-stacks, 2^nd^ to 4^th^ column: single iSIM slices. Second row contains zoomed images of the region in the white box in the top row. White arrows point to F-actin structures protruding deep into the apoptotic cells. Scale bar, 10 μm. **b,** ISIM images of J774 macrophages engulfing DAAM-particles (*E*_y_ 0.3 kPa) functionalized with phosphatidylserine (PS), and TAMRA-Cadaverine for visualization, and immunostained to reveal the exposed surface. Cells were stained for F-actin with AF488 phalloidin. Images were acquired 30 minutes after feeding DAAM-particles. Second row contains zoomed images of the region in the white box in the top row. The white arrow indicates F-actin structures protruding deep into particles and inflicting structural damage.

To establish whether PS alone was sufficient to induce such dynamics, we employed deformable acrylamide co-acrylic acid microparticles (DAAM-particles). These hydrogel particles were functionalized with PS and have low mechanical rigidity (Young’s modulus *E*_y_ 0.3 kPa) and size (∼ 10 μm) similar to apoptotic cells (Lam et al., 2007; Van der Meeren et al., 2020; Vorselen et al., 2020a). J774 macrophages also formed long F-actin-rich protrusions into these artificial targets, which appeared to function as battering rams, leading to disruption of target integrity (Fig. 1B, white arrow). In some cases, this wedging activity could even lead to complete physical breakage and subsequent partial target uptake of the covalently crosslinked, and therefore presumably unbreakable, particles (Fig. 1B). To better visualize the morphology of individual phagocytic cups, we prevented piecemeal uptake of apoptotic cells by fixing them briefly with methanol before feeding them to macrophages. Although J774 macrophages still formed long protrusions into fixed apoptotic A20s, this resulted in a dramatic shift towards whole-cell uptake, allowing us to better visualize the morphology of cups in progress (Fig. S1C,D). Apoptotic cells often appeared embedded in the J774 macrophage cytoplasm before cup closure (Fig. S1C, lower row), which was also observed for unfixed apoptotic A20 cells (Fig. 1A, bottom row) and PS-coated DAAM-particles (Fig. 1B, bottom row). This mode of particle uptake has previously been termed “sinking” phagocytosis and has classically been associated with phagocytosis mediated by complement receptor (CR3, integrin α_M_β_2_, Mac-1, CD11b + CD18) (Allen and Aderem, 1996; Kaplan, 1977; Underhill and Goodridge, 2012).

### PS-mediated Phagocytosis Involves a Sunken Cup Morphology

We hypothesized that the ligands present on the target surface might determine phagocytic morphological dynamics. In order to directly test whether the observed dynamics of apoptotic cell uptake and phosphatidylserine-mediated uptake of DAAM-particles might be distinct from immunogenic antibody-triggered phagocytosis solely because of the ligand identity, we fed otherwise identical DAAM-particles (*E*_y_ ∼ 1.4 kPa) functionalized with either Immunoglobulin G (IgG) or PS (Fig. 2A) to J774 macrophages. For both targets, uptake by J774 macrophages correlated strongly and in a dose-dependent fashion with ligand density on the target (Fig. S2A,B). To reveal potential differences in cytoskeletal remodeling during uptake, cells were fixed and stained for F-actin and imaged with instantaneous structured illumination microscopy (iSIM) (Fig. 2B,D). The unique optical properties of DAAM-particles allowed reliable quantitative measurement of F-actin localization over the entire particle surface (Vorselen et al., 2020b) (Fig. 2C,E). The exposed particle surface was immunostained to precisely determine the stage of engulfment, and thereby to reconstruct phagocytic dynamics in time (Vorselen et al., 2021). For PS-mediated phagocytosis this revealed a unique signature, where in early stages of phagocytic cup formation long actin-rich protrusions (“fingers”) were formed around the target. Such protrusions were rarely observed in IgG-mediated phagocytosis (Fig. 2F).

**Figure 2:**
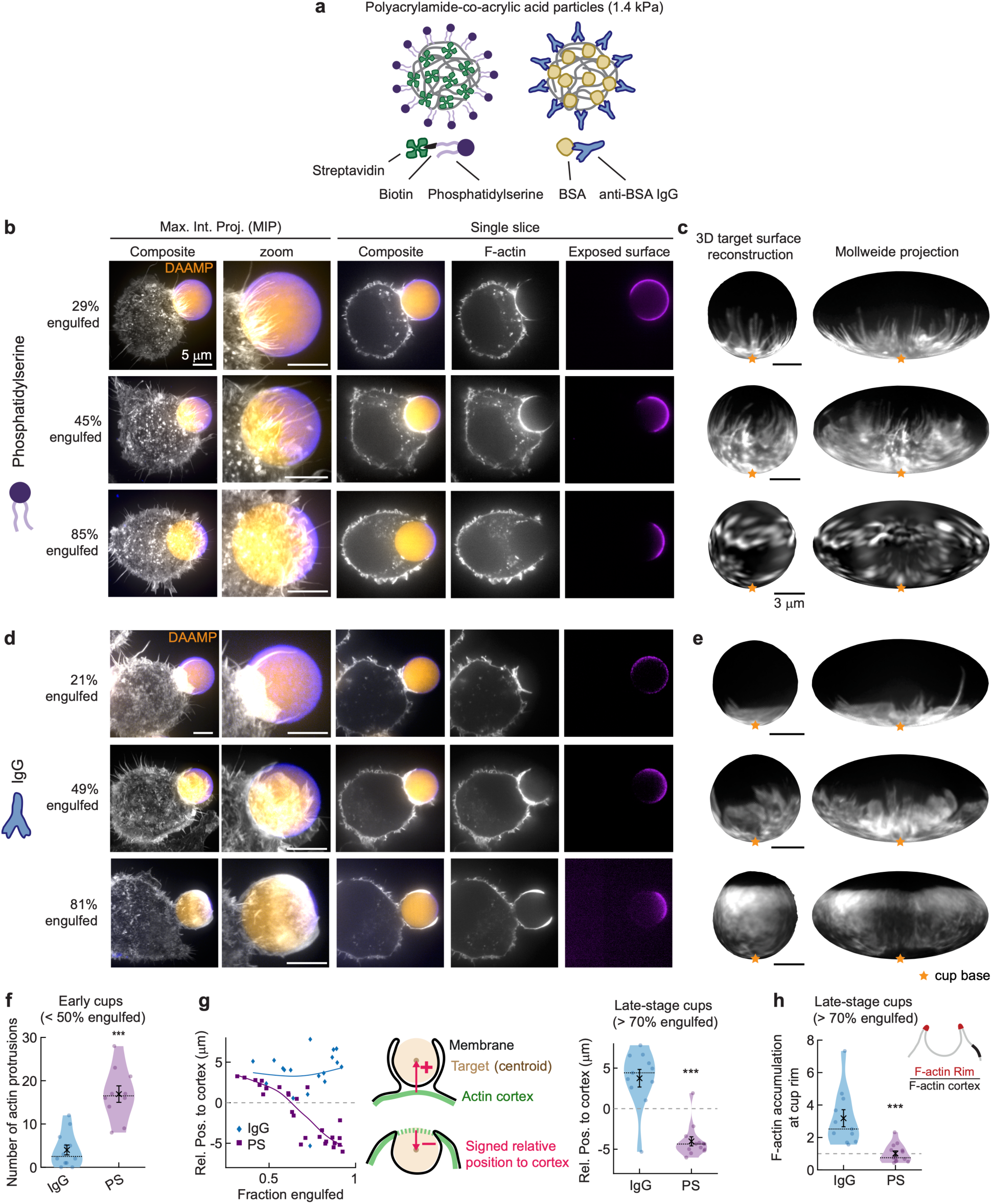
PS-mediated and IgG-mediated phagocytosis have distinct morphological signatures, distinguished by finger-like F-actin protrusions and a sunken cup morphology. **a,** Schematic presentation of the phagocytic targets and functionalization approach. **b,** Superresolution iSIM images of fixed J774 macrophages in various stages of engulfment. Targets were DAAM-particles (*E*_y_ ∼ 1.4 kPa) functionalized with PS, and TAMRA-Cadaverine for visualization, and immunostained to reveal the exposed surface. Cells were stained for F-actin with AF488 phalloidin. Left two columns: composite maximum intensity projections (MIP) of iSIM z-stacks, 3^rd^ to 5^th^ column: single iSIM slices through particle centroid. Scale bar, 5 μm. **c,** F-actin localization visualized on the target surface. Right: Mollweide map projection of entire particle surface. Cups are aligned with the phagocytic axis from bottom to top. Orange stars mark the phagocytic cup base. Scale bars, 3 μm. **d,e,** Similar to **b** and **c**, but for IgG-functionalized DAAM-particles. **f,** Quantification of finger-like actin protrusions in early PS- and IgG-cups (n = 10, 12, respectively). Violin plots show individual phagocytic events (colored markers), mean (black cross), median (dashed line) and standard error of the mean (s.e.m.). *** p = 2×10^-4^. **g,** Left: signed distance of target centroid to (extrapolated) F-actin cortex position. Markers are individual phagocytic events, and lines are cubic smoothing splines showing the trend of the data. Middle: schematic indicating parametrization used for these measurements. Dashed green line indicates extrapolated cortex position. Right: violin plot (styles as in **f**) for late-stage phagocytic cups with n = 11, 13 for PS- and IgG-cups respectively. ***p = 6×10^-4^. **h,** Violin plot (styles as in **f**) showing quantification of F-actin accumulation at the cup rim, measured as illustrated in the insert, for n = 11, 13 PS- and IgG-cups respectively. *** p = 2×10^-4^. Dashed line at 1 corresponds to no F-actin accumulation. Only IgG-mediated cups result in significant actin accumulation (two-sided Wilcoxon signed-rank test for median = 1; IgG: p = 0.001; PS: p = 0.79 not significant). All statistical tests were two-sided Wilcoxon rank-sum tests comparing the IgG- and PS-cups unless otherwise indicated.

In later stages of PS-mediated phagocytosis, the finger-like structures disappeared and the targets often appeared largely embedded in the cytoplasm (Fig. 2B, bottom row). In contrast, IgG-cups had a thin outgrowing pseudopod (Fig. 2D, bottom row). Determination of the relative position of the bead centroid with respect to the actin cortex position for both bead types confirmed no change in bead positioning with phagocytic progression for IgG-cups, and showed “sinking” after 50% engulfment for PS-cups (Fig. 2G). Strikingly, in these late-stage cups there was no significant F-actin accumulation at the cup rim for PS-cups, which is a well-established characteristic for IgG-cups (Fig. 2H, Fig. S2C). DAAM-particles can be used to measure cellular forces (Vorselen et al., 2020b, 2021). Analysis of phagocytic forces revealed that the differences in F-actin localization were accompanied by changes in cellular force exertion patterns (Fig. S3). In IgG-cups, the strongest forces were precisely co-localized with the cup rim where they lead to local target constriction (Fig. S3B). This signature was not observed for PS-cups, where compressive forces were spread throughout the phagocytic cups, and force exertion at the cup rim itself was low (Fig. S3A). Together, these data demonstrate dramatically different morphologies for PS vs. IgG-coated DAAM-particles, where PS-mediated phagocytosis exhibits filopodia-like cellular protrusions, sinking, and weak F-actin polymerization, while IgG cups exhibit broad protrusions, which grow outward and contain strongly polarized F-actin.

### Genome-wide Screens Reveal Specific Regulators of PS- and IgG-mediated Phagocytosis

To identify possible differential genetic requirements for the distinct modes of uptake we observed for PS-vs. IgG-coated DAAM-particles, we designed a genetic screening strategy to identify genes required specifically for uptake of each particle type (Fig. 3A). We developed a method for magnetizing DAAM-particles using iron oxide nanoparticles (see Methods). In parallel screens with each particle type, we incubated the magnetized DAAM-particles with genome-wide knockout libraries of J774 macrophages using conditions in which ∼50% of cells had phagocytosed a particle. We then separated macrophages on a magnetic column into bound and unbound populations, corresponding to macrophages that had or had not phagocytosed a particle, respectively. By sequencing the sgRNA libraries in each population, we identified genes that positively and negatively regulate phagocytosis of each particle type (Fig. 3B-E).

**Figure 3:**
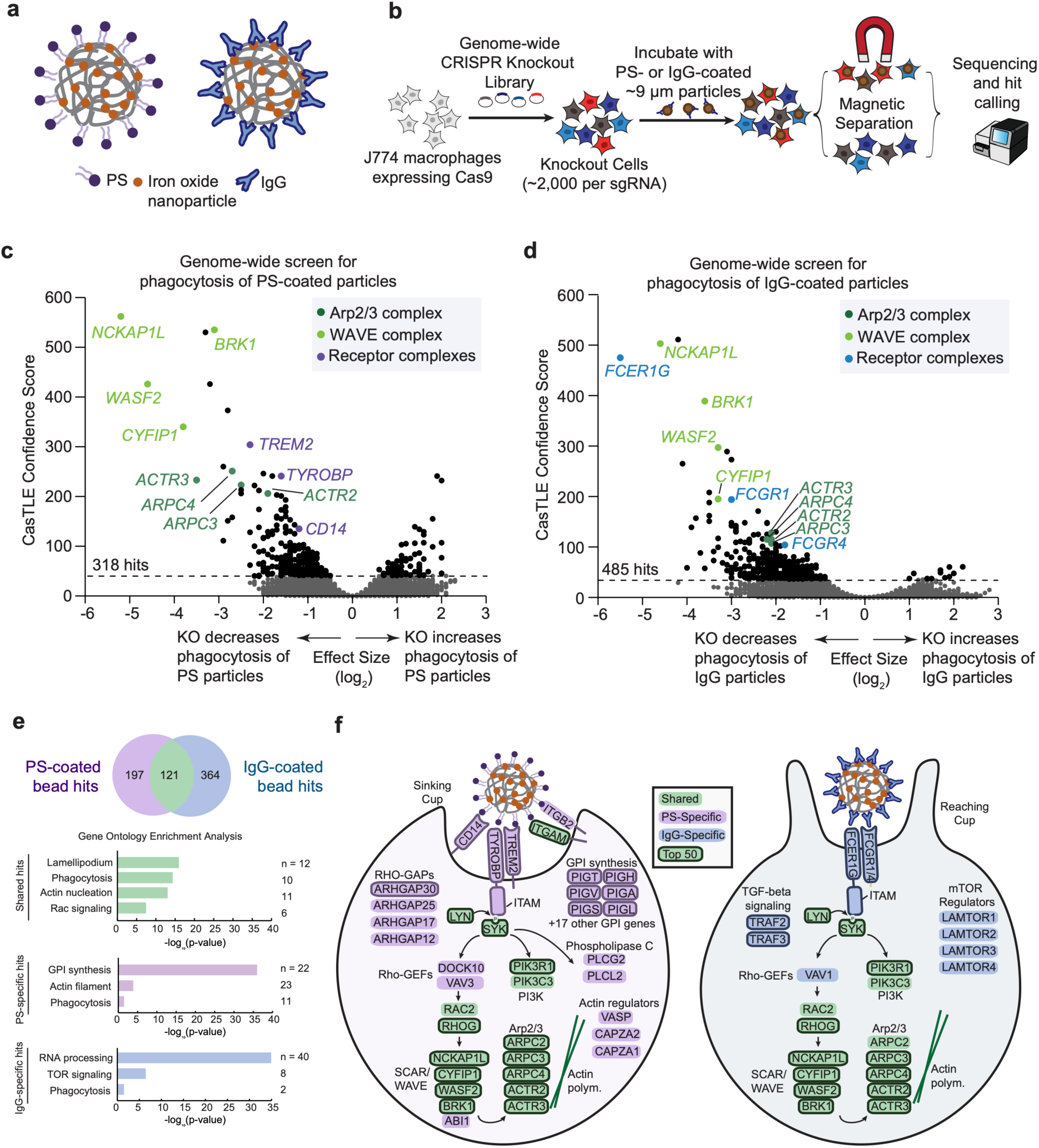
Genome-wide screening reveals shared and specific regulators in PS- and IgG-mediated phagocytosis. **a,** Schematic presentation of magnetized DAAM-particles (*E*_y_ ∼6.5 kPa) synthesized and functionalized for the genome-wide screening approach. **b,** Overview of screening strategy using magnetized DAAM-particles. **c,** Volcano plot of genome-wide screen for uptake of phosphatidylserine-coated DAAM-particles. Dotted line indicates 1% FDR. Total number of hits indicated (including 242 genes whose KO decreased phagocytosis and 76 whose KO increased phagocytosis). **d,** Volcano plot of genome-wide screen for uptake of IgG-coated DAAM-particles. Dotted line indicates 1% FDR. Total number of hits indicated (including 468 genes whose KO decreased phagocytosis and 17 genes whose KO increased phagocytosis). **e,** Top, number of hits (1% FDR) that were shared across screens or unique to each bead type. Bottom, gene set enrichment for shared and unique hits. Selected terms (manually abbreviated for clarity) are shown. n indicates the number of genes in each gene set that are annotated with indicated term. **f,** Schematic of hits from genome-wide screens in **c** and **d**. Top 50 hits in each screen (ranked by casTLE score) are indicated with bold outlines. polym, polymerization.

Given the dramatic structural differences between the phagocytic cups associated with these two different ligands, we were unsure whether to expect significant overlap in cytoskeletal regulators for the two processes. However, we found that members of the Arp2/3 and WAVE complexes that are known actin polymerization regulators during immunogenic phagocytosis (Freeman and Grinstein, 2014; Vorselen et al., 2021) were among the top hits in both screens. Additionally, several genes required for signaling downstream of ITAM-containing phagocytic receptors, including *RHOG, LYN, SYK,* and *PIK3C3,* were identified as top hits in screens with both particle types (Fig. 3F).

The screens also revealed a number of genes and pathways that were unique to each ligand. As expected, several Fc-gamma receptors and the common Fc receptor signaling subunit *FCER1G* were among the strongest hits in the IgG screen, but were not hits in the PS screen. In the PS particle screen, several of the top unique hits likewise encoded cell surface receptors that could be candidates for ligand binding and/or signal transduction during phagocytosis. Many putative receptors have been previously reported for PS-mediated phagocytosis, including BAI-1, TIM-4, Mer, Stabilin 2, CD300, Rage, and integrin α_v_β_3/5_ (Elliott et al., 2017; Gheibi Hayat et al., 2019; Miyanishi et al., 2007; Park and Kim, 2017; Park et al., 2007). Because of the apparent redundancy between receptors, and because of low expression levels of some of these receptors in macrophages (Hsiao et al., 2019), the identities of the critical receptors for PS-mediated phagocytosis in macrophages are still unclear. Surprisingly, our unbiased genome-wide screening approach suggested TREM2 (together with its coreceptor TYROBP), CD14, and integrin subunits ITGAM and ITGB2 as possible receptors for uptake of PS-coated targets. TREM2 is a phagocytic receptor for lipoproteins and neuronal debris, which is highly expressed in microglia and widely studied for its relation to neurodegenerative diseases (Deczkowska et al., 2020; Jay et al., 2017; Yeh et al., 2017). CD14 has known roles in phagocytosis of bacteria through its interaction with lipopolysaccharide (LPS) (Kagan, 2017; Wright et al., 1990). It also has a role in apoptotic-cell uptake (Devitt et al., 1998), potentially involving recognition of ICAM3 or phosphoinositide binding (Kim et al., 2022; Moffatt et al., 1999). Finally, *ITGAM* and *ITGB2* encode the two subunits of the heterodimeric integrin α_M_β_2_, which is the complement receptor, a well-known phagocytic receptor, but different from the integrins (α_v_β_3/5_) previously implicated in PS-dependent phagocytosis (Freeman and Grinstein, 2014).

In addition, many other genes not previously implicated in phagocytosis were identified as unique hits with each particle type. Strikingly, 23 genes required for the synthesis of glycosylphosphatidylinositol (GPI) and its attachment to proteins were among the unique hits in the PS screen whose knockout decreased phagocytic efficiency, corresponding to a large majority of the genes in this pathway (Kinoshita, 2020). In addition, several Rho-GAPs (*ARHGAP30, ARHGAP25, ARHGAP17, ARHGAP12)* and several actin regulators (*VASP, CAPZA1/2*) were identified as unique hits whose knockout decreased efficiency of uptake for PS-coated but not IgG-coated targets. The protease ADAM17, which has been shown to cleave TREM2 and reduce TREM2 levels at the cell surface (Feuerbach et al., 2017), was identified as a negative regulator in the PS screen; that is, its knockout caused an increase in phagocytic uptake. Previously, the guanine exchange factor (GEF) DOCK1 (DOCK180) was established as the key activator of Rac-1 and downstream cytoskeletal remodeling in the context of PS-driven phagocytosis (Gumienny et al., 2001; Penberthy and Ravichandran, 2016). Interestingly, our screen identified several other GEFs, including *VAV3* and *DOCK10*, which may serve as alternative activators of RHO GTPases. In the IgG screen, several members of the LAMTOR complex and the signaling factors *TRAF2* and *TRAF3* were identified as unique hits whose knockout decreased phagocytic efficiency. Intriguingly, knockout of *TREM2* and *TYROBP* caused an increase in phagocytic efficiency for IgG-coated targets, the opposite of their knockout phenotype for PS-coated targets. Overall, these screens recovered a large number of genes previously shown to be required for phagocytosis as well as many genes not previously linked to phagocytosis. Importantly, many of the genes identified as hits only in one screen corresponded to the expected substrate-specific receptors for the ligand presented by each bead type, confirming the biological specificity of the two different screens. The additional unique hits in each screen thus represent novel candidate substrate-specific regulators that may be worthy of further exploration.

### The GPI Anchor Synthesis Pathway is Crucial for PS-Mediated Phagocytosis Through the Involvement of CD14

The single most enriched category of genes in the genome-wide screen for factors whose knockout decreased the efficiency of PS-mediated phagocytosis were genes involved in synthesis and attachment of glycosylphosphatidylinositol (GPI), a lipid anchor for many cell-surface proteins (Fig. 4A) (Kinoshita, 2020). We established a CRISPR knockout line of *PIGX*, which encodes an essential protein for mannose transfer in GPI-anchor precursors during GPI-anchor biosynthesis (Ashida et al., 2005), in J774 macrophages and validated that knockout of *PIGX* specifically inhibited PS-dependent phagocytosis but had no effect on IgG-dependent phagocytosis (Fig. 4B, Fig. S4A,B).

**Figure 4:**
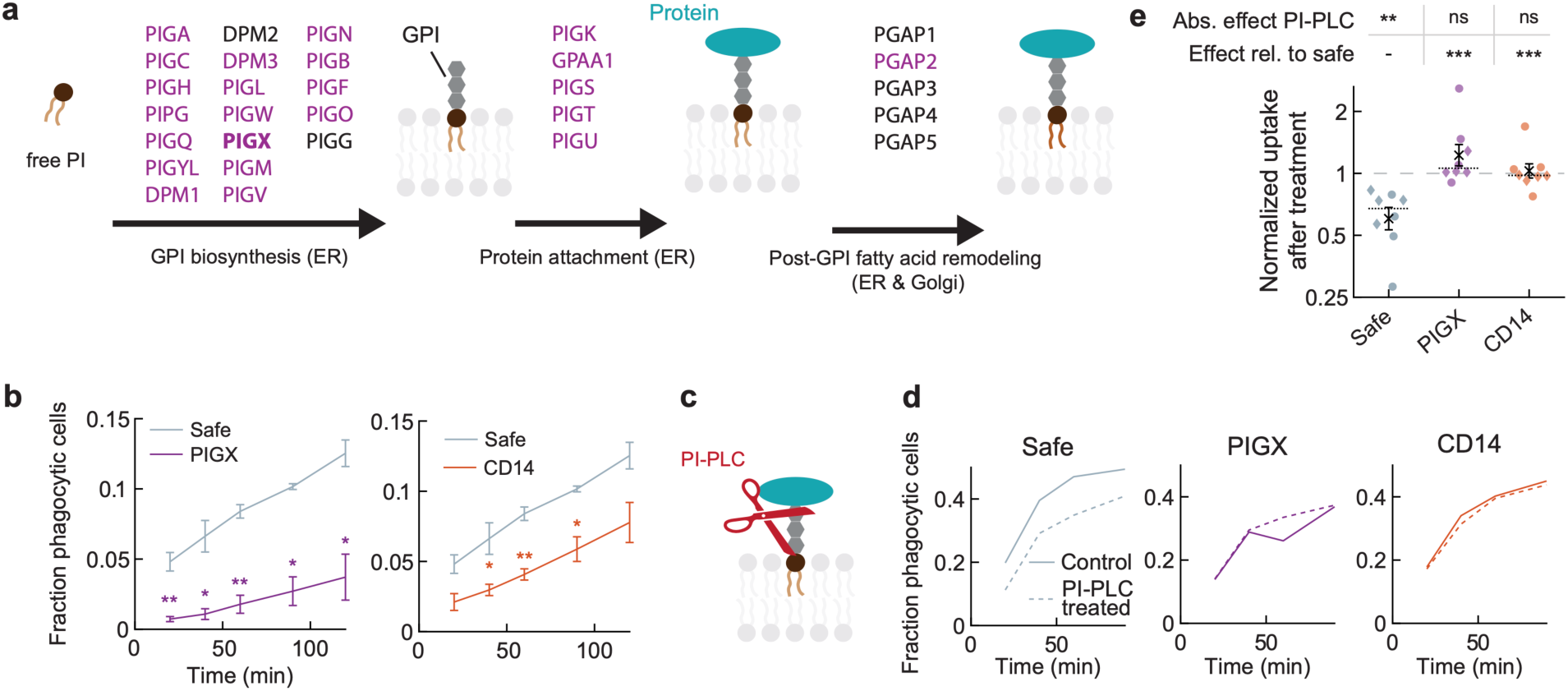
The GPI-synthesis pathway is crucial to PS-mediated phagocytosis through the specific involvement of CD14. **a,** Schematic of the GPI-synthesis pathway with hits in various steps of this pathway (all PS-specific) indicated (ER: endoplasmic reticulum). In bold: PIGX, for which a CRISPR KO line was established. **b,** Phagocytic assay with J774 macrophage PIGX KO and control (safe guide) cells carried out in solution. Error bars indicate standard deviation (s.d.) of three replicate experiments. In total, at least 4×10^5^ cells were analyzed for both cell lines. Two-sided t-tests were used to compare PIGX KO cells with safe guide KO control cells at each timepoint, with significance levels: p < 0.05*; p < 0.01**. **c,** Schematic presentation of the functioning of enzymatic treatment of phosphoinositide-specific phospholipase C (PI-PLC), which specifically cleaves GPI-anchored proteins from the membrane. **d,** Time course showing effect of PI-PLC treatment on phagocytic uptake efficiency by adherent J774 macrophages. A single representative experiment is shown. **e,** Relative uptake of 1 u/mL PI-PLC treated and untreated cells. Individual markers indicate separate experiments for which phagocytic efficiency was evaluated after 20, 40, 60 or 90 minutes (colored markers). Circles and diamonds indicate results from assays carried out with adherent cells (data from **d**) and suspension cells, respectively. Mean (black cross), median (dashed line) and standard error of the mean (s.e.m.: error bars) are indicated. Dashed line at 1 corresponds to no observable treatment effect. Two-sided Wilcoxon signed rank tests indicate treatment effect (*i.e*. median ratio deviating from 1) and two-sided Wilcoxon rank-sum tests compare treatment effects with safe guide KO cells, both with significance levels: p < 0.01**; p < 0.001***.

We considered two broad categories of mechanisms by which the GPI anchor biosynthetic pathway could regulate PS-dependent phagocytosis. First, GPI-anchored proteins could contribute to phagocytosis via biophysical mechanisms unrelated to the functions of the individual proteins anchored to the membrane by GPI. Indeed, GPI anchors have previously been suggested to regulate the formation of plasma membrane lipid nanodomains and receptor nanoclustering (Baranov et al., 2019). Second, the GPI anchor biosynthesis pathway may be required simply for the function of one or more GPI-anchored proteins required for PS-mediated phagocytosis. Relevant to this second possibility, only one of the strong PS-specific hits identified in our screens, CD14, was a GPI-anchored protein. By establishing a CRISPR knockout line of *CD14* in J774 macrophages, we validated the screen result that *CD14* knockout specifically inhibited PS-dependent phagocytosis (Fig. 4B, Fig. S4D). Thus, to experimentally distinguish between these two broad mechanistic possibilities, we sought to evaluate whether defective PS-dependent phagocytosis observed upon loss of GPI-anchored protein synthesis might be primarily due to improper processing of CD14, which we observed in the PIGX knockout (Fig. S4E). We surmised that if GPI anchors are required for PS-mediated phagocytosis solely through supporting CD14 function, acute enzymatic elimination of GPI anchored proteins would inhibit phagocytosis by control macrophages but would have no effect on CD14- or GPI-anchor-deficient macrophages. To this end, we incubated cells with phosphatidylinositol-specific phospholipase C (PI-PLC), which enzymatically cleaved ∼80% of GPI-anchored proteins from the cell surface (Fig. 4C, Fig. S4F). As expected, PI-PLC treatment led to a significant decrease of uptake (∼30%) of PS-coated DAAM-particles by safe control macrophages, but had no further effect on *PIGX* KO cells, which are already deficient in GPI-APs and impaired in phagocytosis (Fig. 4D,E). Importantly, PI-PLC treatment also had no further effect on *CD14* KO cells, which lack *CD14*, but still expose many other GPI-anchored proteins on their surface. These results strongly suggest that the involvement of the GPI-synthesis pathway in PS-mediated phagocytosis is primarily related to the requirement for *CD14* for efficient uptake.

### TREM2, CD14, and ITGAM Contribute Distinctly to Phagocytic Cup Formation

As mentioned above, several of the top differential hits between PS- and IgG-functionalized targets were plasma membrane-localized proteins (Fig. 5A). As expected, this included FCER1G and FCGR1/4, subunits of the Fc receptor (FcR) for IgG-coated targets. For PS-coated targets, TREM2, CD14 and integrin α_M_β_2_ were the most essential cell surface proteins. In individual assays, we confirmed that genetic deletion of these putative PS-receptors, as well as the TREM2 interaction partner TYROBP, inhibited phagocytosis using knockout J774 macrophages of these genes (Fig. 5B, Fig. S4C). To test whether TREM2 and CD14 were able to directly bind to PS, we used protein-lipid overlay assays using recombinant ectodomains of TREM2 and CD14. Consistent with previous reports (Cannon et al., 2012; Wang et al., 2015), this revealed that TREM2 is able to directly bind PS, and we additionally observed binding of TREM2 to multiple phosphorylated phosphatidylinositols (Fig. 5C). In contrast, CD14 was not observed to bind PS directly in this assay.

**Figure 5:**
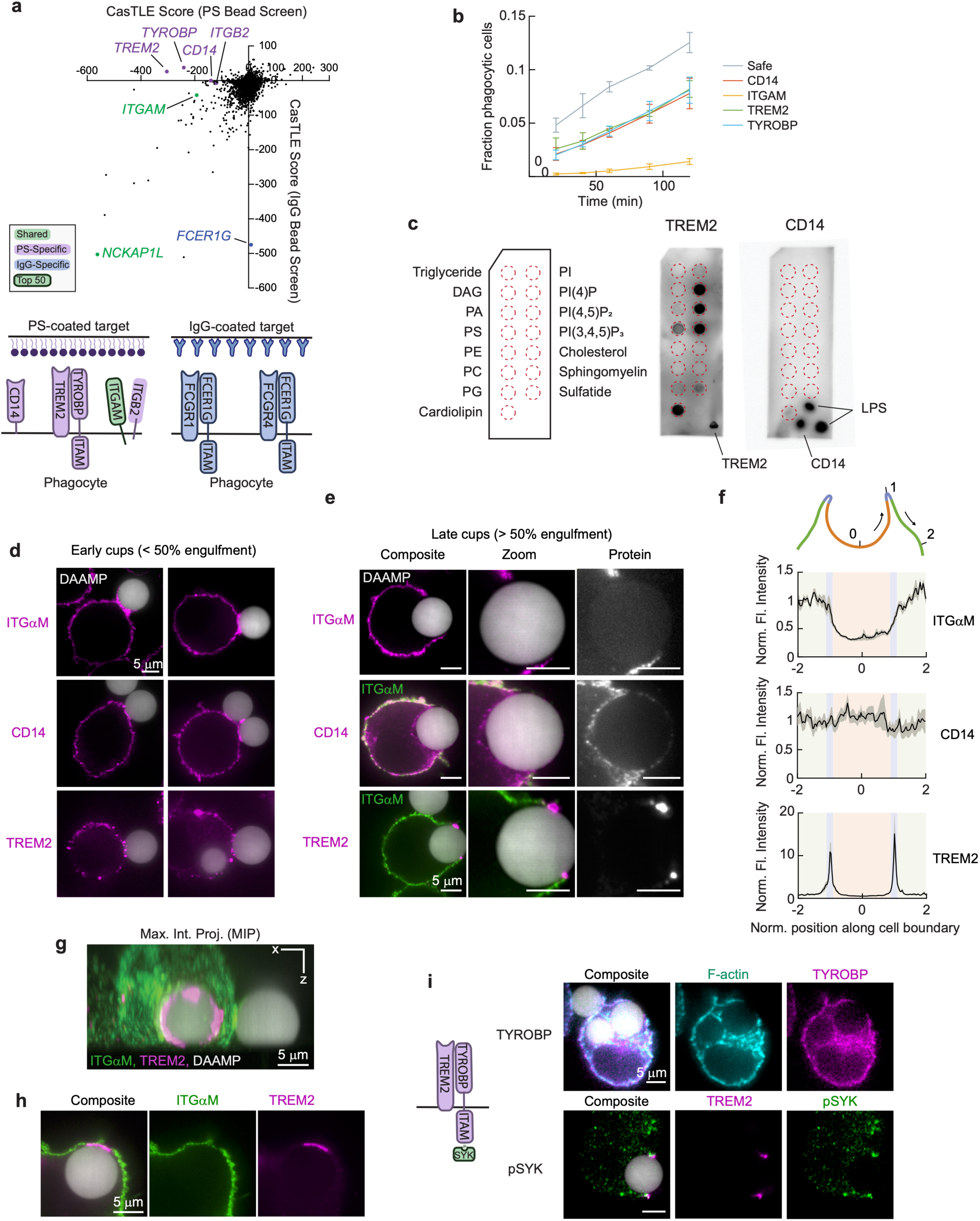
TREM2, CD14 and integrin α_M_ contribute distinctly to PS-mediated phagocytic cup formation. **a,** Comparison of receptor hits in genome-wide screens for regulators of phagocytosis of PS-vs IgG-coated DAAM-particles. Bottom, schematic summary of the putative receptors for PS (left) and IgG (right). **b,** Phagocytic efficiency for control (safe guide) cells, and CRISPR KO lines of putative PS receptors carried out in solution. Error bars indicate standard deviation (s.d.) of three replicate experiments. In total, at least 4×10^5^ cells were analyzed for each cell line. **c,** Protein-lipid overlay assay with 15 different membrane lipids. Left; schematic indicating location of lipid species. Right; binding of recombinant murine TREM2 and CD14 containing Fc domains, detected using a HRP-bound antibody. Additional spots containing positive controls TREM2, CD14 and lipopolysaccharide (LPS) are also indicated. **d,e,** iSIM images (slices) of fixed J774 macrophages during engulfment of 6.5 kPa DAAM-particles functionalized with PS, and TAMRA-Cadaverine for visualization. Putative PS-receptors were immunostained. **d,** Early-stage cups showing receptor accumulation at the cell-bead interface. **e,** Late-stage cups showing distinct localization profiles of ITGαM, CD14 and TREM2. 2^nd^ and 3^rd^ row were counterstained with ITGαM (green) to visualize the cell shape. **f,** Quantification of receptor localization in and around the phagocytic cup. Colored areas and positions correspond to similarly colored sections indicated in the schematic. Fluorescence intensity was normalized to signal at the membrane outside of the phagocytic cups. **g,** Maximum intensity projection in XZ showing TREM2 accumulation in a ring along the entire rim of the phagocytic cup. **h,** Representative image of TREM2 localization after completion of engulfment at the putative site of cup closure. **i,** Representative images of TREM2 interaction partner TYROBP and phosphorylated downstream signaling molecule pSYK in late-stage phagocytic cups.

To gain further insight into possible roles of TREM2, CD14 and ITGAM, we used immunofluorescence staining and super-resolution iSIM imaging to determine whether these proteins localize to phagocytic cups (Fig. 5D,E). During initial contact formation in the early stages of phagocytosis, each of these proteins was found to be enriched at the cell-target interface (Fig. 5D). Strikingly, in later stages of phagocytosis TREM2, CD14 and ITGAM each showed unique localization patterns (Fig. 5E), which we quantified throughout the phagocytic cup (Fig. 5F). CD14 was found throughout the cell-target interface throughout all stages of phagocytic cup development. By contrast, ITGAM localized primarily to early-stage phagocytic cups and to rare outgrowing cups (Fig. S5A), but was largely absent from late-stage cups (Fig. 5E,F). Most strikingly, TREM2 accumulated precisely at the phagocytic cup rim, with a large 10-20 fold enrichment over the concentration measured elsewhere on the cell membrane. We further confirmed this localization by volumetric imaging, and observed that TREM2 exhibited a ring-like localization pattern at the rim of late-stage phagocytic cups (Fig. 5G). In cups that had presumably been closed recently, TREM2 was enriched in a patch corresponding to the apparent site of cup closure (Fig. 5H). At these sites of high TREM2 accumulation, amounts of ITGAM were reduced (Fig. S5B). Given the striking localization of TREM2 at the rim of late-stage phagocytic cups, we also evaluated the localization of TREM2 interaction partner TYROBP, which revealed a uniform staining of the plasma membrane without specific accumulation in cups (Fig. 5I). However, downstream signaling molecule phospho-SYK (tyrosine 342) was highly enriched together with TREM2 in the cup rim, indicative of active TREM2 signaling at these sites (Fig. 5I). The divergent localization patterns of CD14, ITGAM, and TREM2 at the phagocytic cup suggested these cell surface factors may perform distinct functions during PS-dependent phagocytosis.

### Loss of TREM2 and CD14 Results in Aberrant Phagocytic Cup Formation

To investigate the individual contributions of CD14, TREM2 and integrin α_M_β_2_ to the morphology of PS-driven phagocytic cups, we fixed control and knockout cells during uptake of PS-coated targets and imaged their F-actin distributions. No clear deviations from normal cup formation were observed in ITGAM-deficient macrophages. In contrast, upon loss of TREM2, the long F-actin-rich protrusions that we previously observed in early PS cups were strongly reduced in numbers (Fig. 6A), suggesting a role for TREM2 in formation of these finger-like actin structures. In further support of such a function, using immunofluorescence we observed TREM2 at the tips of protrusions both during uptake of PS-coated DAAM-particles and during uptake of apoptotic A20 cells (Fig. 6B, Fig. S5C). ISIM microscopy also revealed an effect of loss of CD14 during phagocytic cup formation. Whereas in normal PS cups there is little F-actin accumulation compared to the F-actin levels in the cell cortex, loss of CD14 resulted in atypical patches of high-intensity F-actin within the phagocytic cup (Fig. 6C). Together, these results indicate that CD14 and TREM2 are required for the normal progression of PS-cup formation.

**Figure 6:**
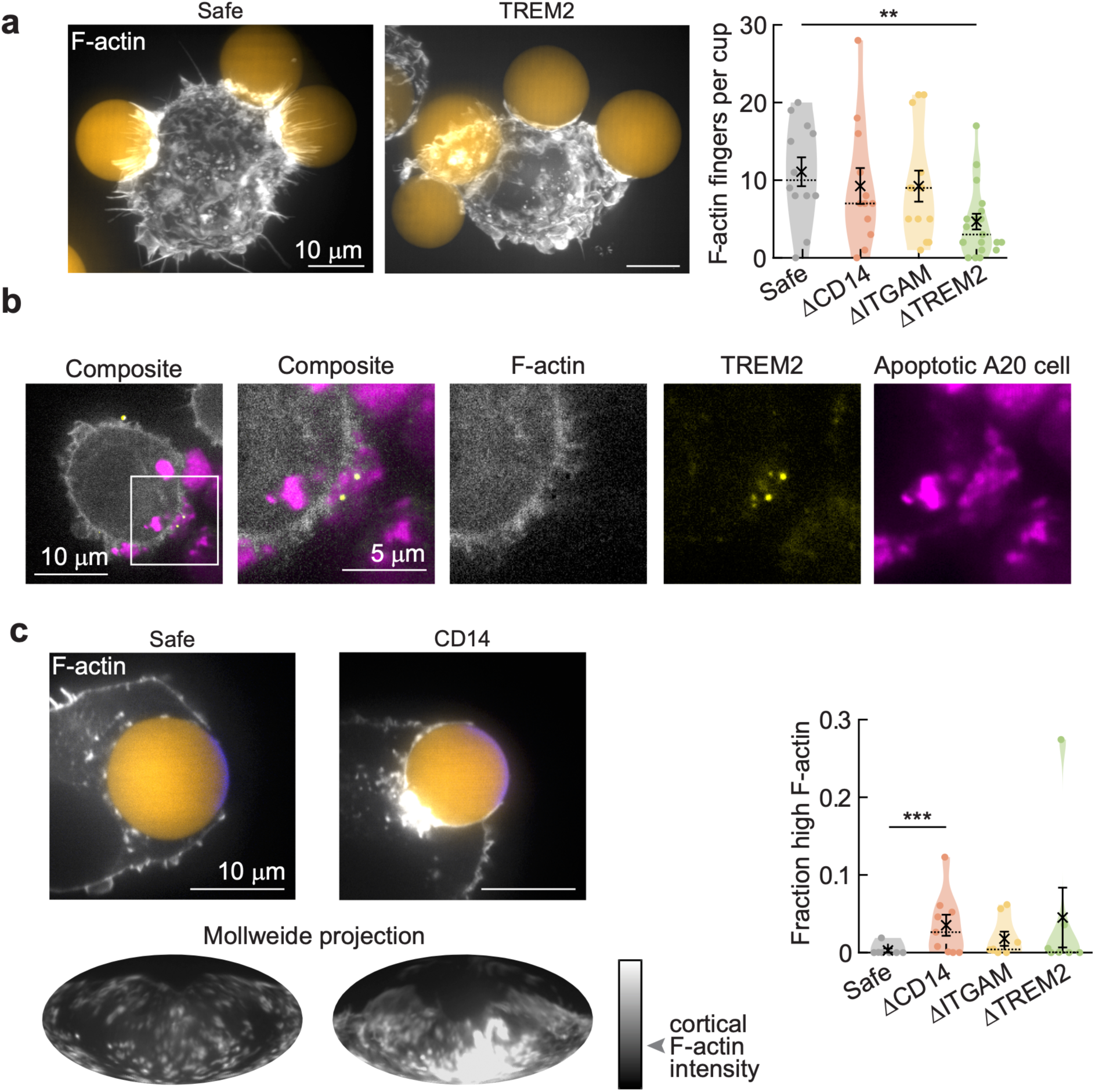
Loss of TREM2 and CD14 results in aberrant cup shaping dynamics. **a,** Left, representative iSIM images (max. Intensity projections) of phalloidin-stained fixed J774 macrophages, either control (safe guide) or CRISPR KO, during engulfment of 6.5 kPa DAAM-particles functionalized with PS, and TAMRA-Cadaverine for visualization. Right, quantification (manual counting) of F-actin fingers (n = 10-15 phagocytic cups per cell line). Individual phagocytic cups (colored markers), mean (black cross) and median (dashed line) and standard error of the mean (s.e.m.) are shown. ** p = 0.008. **b,** iSIM images (single slices) of fixed J774 macrophages, which were transduced with LifeAct-EGFP, interacting with apoptotic A20 cells, which were treated with 5 μM staurosporine for 20 h and stained with CellTrace Far Red. Immunostaining of TREM2 reveals localization at tips of F-actin protrusions during uptake of apoptotic A20 cells. **c,** Similar to a, with single image slices (top) and Mollweide projections normalized to the cortical F-actin intensity (bottom) shown. Right, quantification of the fraction of high F-actin (> 3-fold cortical F-actin intensity). Cortical F-actin intensity was determined in the z-slice through the bead center at a membrane section opposite to the site of bead engagement. *** p = 9×10^-4^. All statistical tests were two-sided Wilcoxon rank-sum tests comparing the distribution median to the control (safe) cells.

### Loss of TREM2 Results in Deficiency of Apoptotic Cell Uptake in Zebrafish Embryos

To test if TREM2 deficiency affects apoptotic cell uptake in an *in vivo* setting, we used a zebrafish developmental model (Herbomel et al., 2001). Significant neuronal remodeling occurs during early zebrafish embryonic development, which includes apoptosis of neurons and engulfment of these apoptotic cells by microglia, the macrophages of the brain (Casano et al., 2016; Peri and Nüsslein-Volhard, 2008). We generated mutant animals (CRISPants) lacking Trem2 by microinjection of Cas9 protein and guide RNAs targeted to the zebrafish homolog, *trem2,* in *Tg(mpeg1:mCherry)* embryos, which express a fluorescent reporter for microglia. Apoptotic cells in *trem2* mutant animals or uninjected control animals were labeled with acridine orange (AO) at 4 days post-fertilization (Fig. 7). We found that a smaller percentage of microglia in *trem2* mutant animals were AO+, indicating a requirement for Trem2 during apoptotic cell engulfment *in vivo* (Fig. 7).

**Figure 7:**
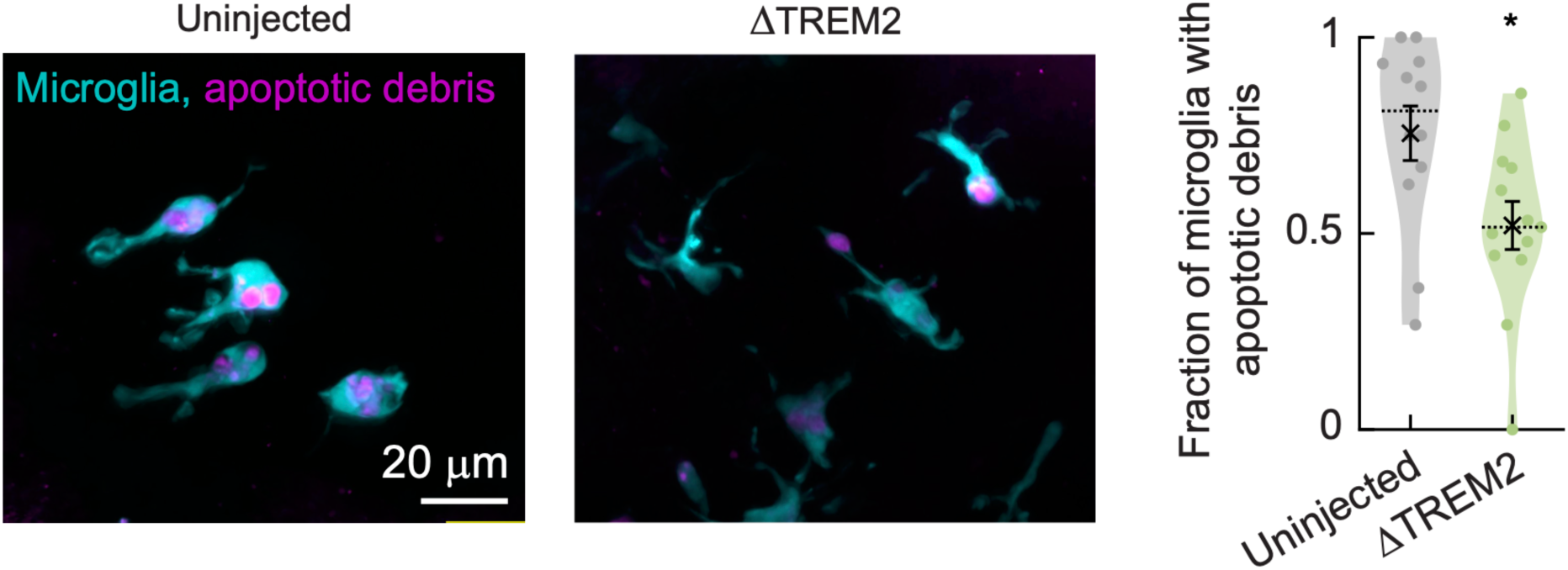
Loss of TREM2 leads to reduced apoptotic cell clearance *in vivo*. Zebrafish embryos were microinjected at the 1-cell stage with Cas9 protein and 3 gRNAs targeted to exon 2 of *trem2.* Uninjected animals of the same clutch were used as controls. Apoptotic debris was stained with acridine orange. Left, representative confocal images (z-stacks). Right, quantification of microglia with internalized cell bodies (n = 12,13 for uninjected and *trem2* mutant fish respectively). Individual fractions per fish (colored markers), mean (black cross) and median (dashed line) and standard error of the mean (s.e.m.) are shown. p = 0.02 determined by a two-sided Wilcoxon rank-sum test comparing median fractions.

## Discussion

Macrophages fail to phagocytose harmful extracellular material in a wide range of diseases, including cancer, atherosclerosis and neurodegeneration, and restoration of proper phagocytosis has thus emerged as a possible therapeutic strategy (Feng et al., 2019; Janda et al., 2018; Kojima et al., 2016). One challenge in the design of phagocytosis-modulating therapies is the need to not only drive engulfment of a particular target, but to ensure that macrophages respond appropriately to the nature of the engulfed target. As an example of this challenge, in cancer treatment, tumor-targeting antibodies such as rituximab are used to drive the phagocytosis of cancer cells, but the clinical efficacy of these antibodies is limited by the induction of an immunosuppressive response by the macrophage following phagocytosis of live or dead cancer cells (Su et al., 2018). Here, we combined a comparative full-genome screening approach with high-resolution imaging to define the distinct molecular mechanisms and morphological features of two forms of phagocytosis, driven by antibodies or phosphatidylserine, that are known to produce immunogenic and non-immunogenic macrophage responses under most conditions. We identify both remarkable convergence in the core actin signaling modules required for both forms of phagocytosis and pronounced morphological and regulatory differences underlying these distinct uptake pathways. These findings advance our understanding of the fundamental biology of phagocytosis and serve as a broadly useful resource that will support efforts to identify strategies for modulating phagocytosis in a defined manner.

The identification of TREM2 as a key regulator of PS-mediated phagocytosis in macrophages is of particular interest because of the broad implication of this gene in neurodegenerative disease and, more recently, in tumorigenesis (Binnewies et al., 2021; Deczkowska et al., 2020; Molgora et al., 2020; Yeh et al., 2017). Loss of TREM2 has previously been shown to reduce phagocytosis of apoptotic debris, lipoproteins, Aβ-amyloid and bacteria in microglia, but the mechanisms of TREM2 involvement in phagocytosis have remained unclear. As most of our studies were conducted in macrophages rather than microglia, our results suggest that TREM2 may be important in macrophage biology beyond microglia and give multiple new insights into its mode of action. Firstly, observations that TREM2 deficiency in microglia does not affect apoptotic cell uptake in some studies have lent support to a model in which TREM2 is involved in phagocytosis only through indirect roles in proliferation or survival of activated phagocytes (Deczkowska et al., 2020; Wang et al., 2015). Our observations of reduced apoptotic cell uptake on a per microglia basis *in vivo*, combined with our findings that TREM2 is required for PS-dependent phagocytosis by macrophages in vitro, localizes to PS-dependent phagocytic cups, and binds PS directly, however, provide strong evidence for direct involvement of TREM2 in phagocytosis. Secondly, we establish that TREM2 clusters at the cup rim, where it actively signals and persists as a micron-scale patch after cup closure. For many immune receptors, cluster formation at the nano- to micron-scale is essential for the activation of downstream signaling (Li and Yu, 2021; van der Merwe and Dushek, 2011). Combined with previous studies indicating that crosslinking of TREM2 increases downstream pSYK activity and phagocytosis (Schlepckow et al., 2020; Takahashi et al., 2005), our results implicate that cluster formation is also essential for TREM2 signaling. Thirdly, if and how TREM2 cooperates with other phagocytic receptors has remained unclear. We show that TREM2 works in concert with CD14 and integrin α_M_β_2_ during clearance of PS-exposing targets. Lastly, the dependence on TREM2 for the formation of F-actin protrusions in early cups, and its accumulation and signaling at the rim of sunken cups, gives insight into how TREM2 affects F-actin remodeling during phagocytic cup formation. Although sinking phagocytosis has typically been connected to integrin α_M_β_2_ (complement receptor) (Allen and Aderem, 1996; Kaplan, 1977; Underhill and Goodridge, 2012), this view has recently been challenged (Jaumouillé et al., 2019; Walbaum et al., 2021). Our observations of TREM2 localization at the rim of sunken cups (Fig. 5e-g) and aberrant F-actin organization in such cups in CD14-deficient cells (Fig. 6c), suggest that sinking phagocytosis is likely not solely dependent on complement, but instead, on the interplay between multiple receptors, including TREM2 and CD14.

We also establish CD14 and integrin α_M_β_2_ as important regulators of PS-mediated phagocytosis. Integrin α_M_β_2_ is best known for its ability to bind the complement component C3b, but it is a promiscuous receptor, especially after inside-out activation by cytoplasmic signals that increase affinity of the integrin extracellular domain for a broad spectrum of ligands (Lamers et al., 2021). Upon activation, integrin α_M_β_2_ may function as a molecular clutch, which transmits the cytoskeletal forces that drive target engulfment (Jaumouillé et al., 2019). CD14, on the other hand, has a known role in apoptotic cell uptake by macrophages (Devitt et al., 1998), but our work sheds light on its mechanistic role, which has long remained elusive. Despite being required for efferocytosis, CD14 does not appear to bind PS directly (Devitt et al., 1998; Thomas et al., 2013), and the possibility that it binds other ligands on apoptotic cells has been proposed (Devitt et al., 1998; Thomas et al., 2013). Our observations that CD14 is required for phagocytosis of microspheres whose uptake is wholly dependent on PS helps clarify that CD14 is indeed required for PS-dependent phagocytosis despite not detectably binding PS. Further studies are required to dissect the mechanistic basis by which CD14 contributes to PS-dependent signaling, but our observation of altered actin dynamics in CD14-KO macrophages suggests that CD14 may regulate actin dynamics during PS-dependent phagocytosis. Interestingly, the shared contributions of CD14 and integrin α_M_β_2_ we observe in PS-mediated uptake, has previously been observed in uptake of the bacterium *Borrelia burgdorferi (Hawley et al., 2012)*, and hence cooperation between CD14 and integrin α_M_β_2_ may be general to diverse targets of phagocytosis.

We identified the formation of long F-actin-rich protrusions as a characteristic morphological feature of PS-dependent phagocytosis. The general involvement of such F-actin “fingers’’ in PS-mediated phagocytosis is implied by their formation during engulfment of apoptotic A20 cells as well as PS-coated DAAM particles of multiple rigidities. These structures protrude several microns into apoptotic cells and softer particles, whereas on higher rigidity targets they are likely incapable of doing so, and instead, are deflected and follow the target surface. The capability of these F-actin protrusions to induce structural damage on artificial targets and their frequent coincidence with engulfed fragments of apoptotic cells may relate to our observation that apoptotic cells are frequently engulfed in piecemeal fashion, a form of uptake also known as trogocytosis or nibbling. Trogocytosis is a widespread behavior among immune cells and beyond (Hamieh et al., 2019; Huang et al., 1999; Ralston et al., 2014; Zhao et al., 2022), and has even been observed during phagocytosis of apoptotic cells *in vivo (Hoijman et al., 2021)*. Importantly, our observations diverge from the general model for trogocytosis in that these protrusions appear to act as wedges, breaking up the target from the inside-out, instead of pinching bits off from the target exterior (Zhao et al., 2022). Such a mechanism may be favorable for fragmenting targets and is likely more common, since protrusions have previously been observed during trogocytosis-mediated killing of cancer cells (trogoptosis) by neutrophils (Matlung et al., 2018).

The regulation of sinking phagocytosis, and specifically the role of the cytoskeleton in this process, is unclear. With the exception of a handful of genes including *CAPZA1* and *CAPZA2*, which are uniquely required for PS-mediated phagocytosis, our screen implies largely overlapping involvement of actin binding proteins for both outgrowing and sunken cups. How does an apparently similar set of proteins create such distinct cytoskeletal structures? One possibility is that cup morphology is determined further upstream, such as by differential involvement of Rho GTPases, which are master organizers of actin dynamics. Indeed, activity of distinct Rho GTPases has been implicated in sinking (RhoA) and outgrowing phagocytosis (Rac1, CDC42) (Caron and Hall, 1998), although our screen suggests shared involvement of Rac1, Rac2 and RhoG in both modes of uptake. Nonetheless, we did identify distinct Rho GEFs, which activate Rho GTPases, in PS-mediated phagocytosis (*DOCK10*, *VAV3*) vs. IgG phagocytosis (*VAV1*). Together with specific involvement of multiple GTPase activating proteins (*ARHGAP30*, *ARHGAP25*, *ARHGAP17* and *ARHGAP12*) in PS-mediated phagocytosis, these upstream regulators likely affect the spatial and temporal dynamics of Rho GTPase activity and cytoskeletal remodeling. Elucidating such regulatory differences between different modes of uptake could ultimately pave the way for the rational design of phagocytosis-modulating therapies that drive or repress engulfment of specific targets.

## Methods

### Cell Culture

J774 Cas9 cells (a gift from A. Sil, UCSF) were cultured in DMEM supplemented with 10% FBS (Gibco), which was heat-inactivated at 56°C for 30 minutes, 100 µg ml-1 streptomycin, 100 U ml-1 penicillin, and 2mM glutamine in a humidified tissue culture incubator set to 37 °C and 5% CO_2_. A20 cells were cultured in RPMI-1640 (ATCC, 30-2001) supplemented with 10% FBS (Gibco), 100 µg ml-1 streptomycin, 100 U ml-1 penicillin, 2mM GlutaMAX, and 50 μM 2-mercaptoethanol in a humidified tissue culture incubator set to 37°C and 5% CO_2_.

### Particle Synthesis

DAAM-particles were synthesized using a membrane-emulsification approach as previously described (Vorselen et al., 2020b). Mixtures of acrylamide containing 100 mg/mL acrylic components, 150 mM NaOH, 0.3% (v/v) tetramethylethylenediamine (TEMED) were prepared in 150 mM MOPS buffer (pH 7.4, made from MOPS sodium salt). Acrylic acid mass fraction was kept constant at 10%, whereas crosslinker mass fraction was 0.32%, 0.65% or 2.3%, for 0.3, 1.4 kPa and 6.5 kPa particles, respectively. For synthesis of magnetic DAAM-particles, 5 mg/mL iron oxide nanoparticles (IONPs, Sigma 747467) was added to the acrylamide mixture. The gel mixture was degassed for 15 min and kept under nitrogen atmosphere until emulsification was completed. 1.9 μm tubular hydrophobic Shirasu porous glass (SPG) were sonicated in n-heptane under vacuum, mounted on an internal pressure micro kit extruder (SPG Technology Co.) and immersed into an oil phase (∼125 mL) consisting of hexanes (99%) and 3% (v/v) Span 80 (Sigma-Aldrich, S6760). 10 mL of gel mixture was extruded into the oil phase under nitrogen pressure of approximately 7 kPa, while the oil phase was stirred continuously at 300 rpm. After completion of extrusion, polymerization of DAAM-particles was induced by addition of ∼225 mg 2,2’-Azobisisobutyronitrile (AIBN) (1.5 mg/mL final concentration) and by raising the temperature to 60°C. The polymerization reaction was continued for 3 h after which the temperature was reduced to 40°C and the reaction was continued overnight. Polymerized DAAM-particles were subsequently washed (5× in hexanes, 1× in ethanol), dried under nitrogen flow for approximately 30 min, and resuspended in PBS (137 mM NaCl, 2.7 mM KCl, 8.0 mM Na_2_HPO_4_, 1.47 mM KH_2_PO_4_, pH 7.4) and stored at 4°C.

### Particle Functionalization

DAAM-particles were functionalized as previously described (Vorselen et al., 2020b). In brief, DAAM-particles were diluted to 5% (v/v) concentration and washed twice in activation buffer (100 mM MES, 200 mM NaCl, pH 6.0). They were then incubated, under gentle mixing, for 15 min in activation buffer supplemented with 40 mg/mL 1-ethyl-3-(3-dimethylaminopropyl) carbodiimide, 20 mg/mL N-hydroxysuccinimide (NHS) and 0.1% (v/v) Tween 20. They were spun down (10,000 × g, 1 min) and washed, each time mixing vigorously, 4× in PBS, pH 8 (adjusted with NaOH) with 0.1% Tween 20. Particles were then resuspended with protein solution. Particles were then functionalized with 5 mg/mL BSA (Sigma, A3059) or streptavidin (Neuromics, 2-0203-100, 0.3 mg/mL) in PBS, pH 8 and incubated while rocking for 1 h. Then cadaverine-conjugate was added: either Tetramethylrhodamine Cadaverine (Thermo Fisher Scientific, A1318), FITC Cadaverine (Thermo Fisher Scientific, A10466) or Cyanine5 amine (Lumiprobe, 130C0) or a combination thereof, to a final concentration between 0.1 - 0.5 mM. After 30 min, 100 mM Tris and 100 mM ethanolamine (pH 9) were added to block unreacted NHS groups. DAAM-particles were then spun down (16,000 × g, 2 min) and washed 4× in PBS, pH 7.4 with 0.1% Tween 20. BSA-functionalized DAAM-particles were washed 3× in sterile PBS and opsonized with 0.02 - 0.05 mg/mL rabbit anti-BSA antibody (MP Biomedicals, 0865111) for 1 h at room temperature. Streptavidin-functionalized DAAM-particles were washed in 50% methanol:water (50% MeOH), and resuspended at 10% solids (v/v) in 0.025 mg/mL biotin-phosphatidylserine (Echelon L-31B16) in 50% MeOH. Finally, DAAM-particles were then washed 3× (10,000 × g, 1 min) with PBS and resuspended in sterile PBS.

### Microscopy

3D instantaneous structured illumination (iSIM) imaging was performed using a Visitech iSIM mounted on a Nikon TiE, equipped with a 100× NA 1.49 objective (Nikon). Images were captured using dual ORCA-fusion (Gen III) CMOS cameras (Hamamatsu). The setup was operated using MicroManager 2.0 (Edelstein et al., 2014).

### Staurosporine treatment and apoptotic cell staining

A20 cells were spun down and resuspended at 1 million cells/mL in DMEM + 10% heat-inactivated FBS with 5 μM staurosporine. After 20 h 5-fold excess of medium was added, cells were spun down and washed once in PBS.

### Anhnexin-V staining

Annexin-V Alexa Fluor 647 (Invitrogen, A23204, 1:20) and nuclear dead cell stain SYTOX orange (Invitrogen, S34861, 1:1000) were diluted in Annexin-V binding buffer (10 mM HEPES (pH 7.4), 0.14 M NaCl, and 2.5 mM CaCl_2_). Cells (10^6^ mL^-1^) were stained for 10 minutes at room temperature. Then, they were put on ice, diluted 5× in Annexin-V binding buffer and measured directly by flow cytometry.

### Phagocytosis Assays

For high-resolution microscopy assays DAAM-particles were added to a total volume of 300 μL of DMEM, supplemented with heat-inactivated FBS, and fed to phagocytes in a 12-well plate. The plate was spun at 300 × *g* for 1 min at room temperature. Cells were incubated at 37°C to initiate phagocytosis for the indicated times (see Figure legends). Cells were then fixed with 4% formaldehyde in PBS for 10 min. Any unbound DAAM-particles were removed by 3 washes in PBS. For visualization of the exposed DAAM-particle area, in case of PS DAAM-particles, samples were incubated with rabbit streptavidin antibody for 1 h (Genscript, A00621, 0.1 mg/mL). IgG and anti-streptavidin treated PS DAAM-particles were stained with donkey anti-rabbit-Alexa Fluor-647 antibodies (Invitrogen, A31573, 1:1000 in PBS) for 30 min. Cells were then washed 3×. In case of F-actin staining with phalloidin, cells were permeabilized with 0.1% Triton X-100 in PBS for 3 min, then stained with Alexa Fluor-488 conjugated phalloidin (Invitrogen, A12379, 0.15 μM). Cells were imaged in PBS or VECTASHIELD Antifade Mounting Medium (Vector Laboratories, H-1000). Phagocytic assays with apoptotic A20 cells were performed similarly. After treatment with staurosporine (STS), A20s were incubated in Celltrace Far Red (Invitrogen, C34564, 1:1000) in PBS for 10 minutes. A20 cells were washed twice in PBS before resuspension in DMEM + hiFBS and use in phagocytic assays. Phagocytic efficiency assays were either carried out on plates or in suspension in tubes, as indicated for individual figures. Phagocytic assays in solution were carried out in 1.5 mL L15 + 10% heat-inactivated FBS. Cells were harvested and resuspended in round bottom culture tubes at 3 × 10^6^ cells/mL or 5 × 10^5^ cells/mL, for PS- and IgG-assays respectively. Functionalized DAAM-particles (8.9 μm diameter, 6.5 kPa) were added at a 3:1 ratio to cells, after which cells were incubated while shaking at an angle at 37°C. At the indicated time points, 0.1 mL was taken from the sample tube, and 4% formaldehyde in PBS was added to fix the cells for 10 minutes. Samples were then washed 3× in PBS and external particles were immunostained as described above. Flow cytometry was used to count large numbers of events. Based on scattering properties, DAAM-particle fluorescence and immunostaining of external particles, the fraction of cells with at least a single internalized particle could be determined (Fig. S4A). For plate based assays, 4 × 10^5^ cells were added per well in 6-well plates and FITC/TMR-dual labeled DAAM-particles (8.9 μm diameter, 6.5 kPa) were used. Wells were scraped at the indicated time point, put on ice, and directly measured on the flow cytometer. The dual-labeled particles allow fully internalized particles to be distinguished from external particles, because FITC fluorescence quenches during phagosome acidification, while TMR fluorescence remains constant.

### Immunofluorescence

Cell surface proteins were stained in the absence of, or before, membrane permeabilization. Cells were incubated with primary antibody for 1 h for ITGAM (Abcam, ab8878, 1:100), TREM2 (Invitrogen, MA531267, 1:100) or CD14 (Abcam, ab221678, 1:250) in PBS. After 3 washes with PBS, they were subsequently stained for 30 minutes with appropriate secondary antibodies 1000× diluted in PBS (Goat anti-Rat AF488 (Invitrogen, A11006), Goat-anti-Rabbit AF488 (Invitrogen, A11008) or Donkey-anti-Rabbit AF647 (Invitrogen, A31573). TYROBP and pSYK-staining were performed after permeabilization (see phagocytic assays for details). TYROBP (Abcam, ab283703, 1:100) and pSYK (BD, pY348, 1:200) staining were performed for 1h, after which cells were washed 3×. Finally, cells were stained for 30 minutes with 1000× diluted donkey-anti-rabbit AF647 (Invitrogen, A31573) or goat-anti-mouse AF488 (Invitrogen, A11001), respectively. Immunostaining for flow cytometry assays was performed at 4°C in PBS pH 7.4 supplemented with 5% FBS and 2 mM EDTA. Both primary CD14 (Abcam, ab221678, 1:500) and secondary Goat-anti-Rabbit AF488 (Invitrogen, A11008, 1:500) incubations were 30 minutes, with 3 washes in ice-cold buffer in between. Afterwards, cells were washed once in buffer, resuspended in 5× original volume, and directly measured on a flow cytometer.

### Genome-wide screens

A ten-guide-per-gene CRISPR knockout library (Morgens et al., 2017) was integrated into J774 Cas9 cells and selected with puromycin, as previously described (Kamber et al., 2021). The genome-wide J774 knockout library was expanded in 15 cm dishes and plated so as to achieve 2000× coverage of the library (∼500 million cells; 20 million cells per plate) in two replicates per screen. To initiate phagocytosis, media was aspirated and replaced with media containing DAAM-particles (∼2 beads per cell). Cells were incubated for 60 minutes with DAAM-particles in a tissue culture incubator, then media was aspirated and cells washed 3× with PBS to remove uneaten beads before macrophages were lifted via mechanical scraping. Cells were pelleted by centrifugation at 300 × g for 5 minutes and resuspended in PBS, then were applied to Miltenyi MACS LS columns that were equilibrated in PBS. Unbound cells were collected and combined with three washes of the column with 2 mL PBS. Bound cells were eluted from the column with 3ml PBS. All samples were pelleted at 500 × g for 5 minutes and stored at −20°C before further processing. Genomic DNA was extracted from each sample using a Qiagen Blood Maxi Genomic DNA Kit, and sgRNA-containing loci were amplified via nested PCR that introduced unique 6 bp barcodes for each sample, and sequenced using a custom sequencing primer on an Illumina NextSeq 550. Samples were demultiplexed using bcl2fastq and hits were calculated using CasTLE (Morgens et al., 2016). A 1% FDR threshold was used to define hits in Fig. 3. Complete genome-wide screening results are presented in Tables S1 and S2.

### Generation of cell lines

For generating stable LifeAct-expressing cells, J774 macrophages were infected with lentiviral constructs expressing LifeAct-EGFP along with blasticidin resistance cassette (Addgene #84383) (Padilla-Rodriguez et al., 2018). At 3 d after infection, cells were selected with 10 μg ml^-1^ blasticidin for an additional 3 d, and EGFP-positive cells were further selected by FACS sorting. To generate knockouts, J774 Cas9 cells were infected with lentiviral constructs expressing indicated sgRNAs and selected with puromycin (5 µg ml^-1^) or blasticidin (10 µg ml^-1^) for 2 d, and cultured for at least 3 d before use in assays.

### Gene ontology enrichment analysis

Gene ontology enrichments were calculated using gProfiler (version e104_eg51_p15_2719230) (Reimand et al., 2016) as ordered queries. Manually selected gene ontology terms are shown.

### PI-PLC treatment

Phosphatidylinositol-specific phospholipase C (PI-PLC) treatment was carried out by adding 1 mL L15 (Gibco, 21083027) + 10% heat-inactivated FBS (hiFBS) with PI-PLC (Invitrogen, P6466, 1 u/mL) per well in a 6-well plate. Cells were treated for 1 h, after which they were washed in fresh medium. Phagocytic assays were carried out directly after treatment.

### Protein-lipid overlay assays

Indicated proteins (see below; 0.1 μg in 1 μL), and mixtures of biotin-LPS (Invivogen, tlrl-lpsbiot) and streptavidin (Neuromics, 2-0203-100) in PBS (pH 7.4) were added to membrane lipid strips (Echelon biosciences, P-6002) as positive controls. Biotin-LPS controls contained 0.5 μg of both streptavidin and biotin-LPS (1 μL total) or 0.2 μg of biotin-LPS and 2 μg streptavidin (2 μL total), for the upper left and lower right spots in Fig. 5C, respectively. Strips were dried for 0.5 h and then blocked in PBS + 30 mg mL^-1^ fatty acid-free BSA (blocking buffer) for 1 h at room temperature. Membranes were incubated in the same buffer for 1 h under light agitation with the recombinant murine chimera proteins containing human Fc domains and the extracellular domains of TREM2 (R&D systems, 1729-T2-050, 0.1 mg/mL) or CD14 (R&D systems, 982-CD-100, 1 mg/mL). Strips were washed 3 × (5 minutes agitation each) in blocking buffer with 0.1% (v/v) Tween 20 added. They were then incubated with Goat-anti-human HRP antibody (Jackson, 109-035-0488, 1:10,000), washed 5 × afterwards and detected using chemiluminescence.

### Zebrafish experiments

To generate *trem2* CRISPants, *Tg(mpeg1:mCherry)^gl23Tg^*zebrafish embryos (Ellett et al., 2011) were microinjected at the 1-cell stage with Cas9 protein and 3 gRNAs targeted to exon 2 of *trem2.* Uninjected animals of the same clutch were used as controls. Lysis of individual embryos followed by PCR genotyping and sequencing was used to ensure gRNA editing efficiency. Starting at 24 h post fertilization, embryos were raised in 0.003% 1-phenyl-2-thiourea to prevent pigmentation. Embryos were screened for mCherry+ microglia at 4 days post fertilization and used for subsequent imaging. Embryos were stained with 1 μg/mL acridine orange (AO) for 1 minute, quickly washed, and embedded dorsal side up in 1% low melt agarose for confocal microscopy. Multiple randomly selected fields of view were imaged per fish, and microglia were scored as AO-positive if they had internalized AO-stained cell bodies. The average number of microglia that were AO-positive across all ROIs per fish was recorded. All zebrafish experiments were approved by the Institutional Animal Care and Use Committee at the University of Washington (Protocol: #4439-01).

### 3D microparticle shape reconstruction and fluorescence maps

Image analysis was performed with custom software in Matlab (Vorselen et al., 2020b, 2021). Cubic interpolation was used to calculate the intensity values along lines originating from the particle centroid and crossing the particle edge from volumetric images. Edge coordinates were then directly localized with super-resolution accuracy by fitting a Gaussian to the discrete derivative of these line profiles. F-actin was mapped to the particle surface using a radial maximum intensity projection within 0.5 μm from the particle edge.

### Force calculations

A fast spherical harmonics method implemented in Python package ShElastic was used for force calculations (Vorselen et al., 2020b; Wang et al., 2019). To derive both normal and shear forces, we solve the inverse problem of inferring the traction forces **T** in an iterative process. We start with a trial displacement field ***u***, and during optimization the following cost function is minimized, while the experimentally determined particle shape is always matched exactly:

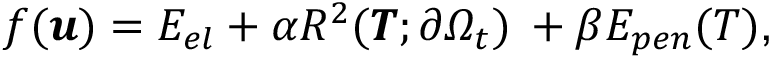

where *αR*^2^(***T***; ∂*𝛺_t_*) are the obtained forces outside of the cell-target contact area, which is obtained from fluorescent F-actin signal, *E_el_* is the elastic energy included to penalize unphysical solutions where higher forces produce the same shape, and *βE_pen_*(*T*) is an anti-aliasing term. The weighing parameter *α* for residual traction outside of the cell-target contact area and *β* for anti-aliasing were both 1. Force measurements were made in the small strain regime (*ε* ≲ 0.1), where particles were assumed to behave linearly elastic. Tractions were evaluated on a 21 × 41 grid and calculated with spherical harmonic coefficients up to *l*_max_ = 20.

## Acknowledgments

This work was supported by an NIH Director’s New Innovator award (1DP2HD084069-01) to M.C.B, R00 HD086271 from the NIH/NICHD to J.P.R., and research support from the Howard Hughes Medical Institute to J.A.T. R.A.K. was supported by a Stanford School of Medicine Dean’s Postdoctoral Fellowship, Jane Coffin Childs Postdoctoral Fellowship, and NIH/NCI K99 Pathway to Independence Award (1K99CA259218-01). D.V. was supported by a CRI Irvington fellowship from the Cancer Research Institute. E. P. was supported by a Washington Research Postdoctoral fellowship. R.L.D.L. was supported by the Stanford Medical Scientist Training Program, National Institutes of Health T32-GM007365 J.P.R. is a Washington Research Foundation Distinguished Investigator.

This article is subject to HHMI’s Open Access to Publications policy. HHMI lab heads have previously granted a nonexclusive CC BY 4.0 license to the public and a sublicensable license to HHMI in their research articles. Pursuant to those licenses, the author-accepted manuscript of this article can be made freely available under a CC BY 4.0 license immediately upon publication.

## Author contributions

Conceptualization: D.V., R.A.K., M.C.B., J.A.T.

Methodology: D.V., R.A.K., R.L.D.L., A.P.vL., E.P.

Formal analysis: D.V., R.A.K.

Investigation: D.V., R.A.K., R.L.D.L., A.P.vL., E.P., M.K.D., S.L.

Writing - original draft: D.V., R.A.K.

Writing - review and editing: R.L.D.L., A.P.vL., E.P., M.K.D., S.L., J.P.R., M.C.B., J.A.T.

Visualization: D.V., R.A.K. Supervision: J.P.R., M.C.B., J.A.T.

Funding acquisition: J.P.R., M.C.B., J.A.T.

## References

Aderem, A., and Underhill, D.M. (1999). Mechanisms of phagocytosis in macrophages. Annu. Rev. Immunol. 17, 593–623.

Allen, L.A., and Aderem, A. (1996). Molecular definition of distinct cytoskeletal structures involved in complement- and Fc receptor-mediated phagocytosis in macrophages. J. Exp. Med. 184, 627–637.

Ashida, H., Hong, Y., Murakami, Y., Shishioh, N., Sugimoto, N., Kim, Y.U., Maeda, Y., and Kinoshita, T. (2005). Mammalian PIG-X and Yeast Pbn1p Are the Essential Components of Glycosylphosphatidylinositol-Mannosyltransferase I. Mol. Biol. Cell https://doi.org/10.1091/mbc.e04-09-0802.

Bäck, M., Yurdagul, A., Jr, Tabas, I., Öörni, K., and Kovanen, P.T. (2019). Inflammation and its resolution in atherosclerosis: mediators and therapeutic opportunities. Nat. Rev. Cardiol. 16, 389–406.

Baranov, M.V., Olea, R.A., and van den Bogaart, G. (2019). Chasing Uptake: Super-Resolution Microscopy in Endocytosis and Phagocytosis. Trends Cell Biol. 29, 727–739.

Binnewies, M., Pollack, J.L., Rudolph, J., Dash, S., Abushawish, M., Lee, T., Jahchan, N.S., Canaday, P., Lu, E., Norng, M., et al. (2021). Targeting TREM2 on tumor-associated macrophages enhances immunotherapy. Cell Rep. 37, 109844.

Boada-Romero, E., Martinez, J., Heckmann, B.L., and Green, D.R. (2020). The clearance of dead cells by efferocytosis. Nat. Rev. Mol. Cell Biol. 21, 398–414.

Cannon, J.P., O’Driscoll, M., and Litman, G.W. (2012). Specific lipid recognition is a general feature of CD300 and TREM molecules. Immunogenetics 64, 39–47.

Caron, E., and Hall, A. (1998). Identification of two distinct mechanisms of phagocytosis controlled by different Rho GTPases. Science 282, 1717–1721.

Casano, A.M., Albert, M., and Peri, F. (2016). Developmental Apoptosis Mediates Entry and Positioning of Microglia in the Zebrafish Brain. Cell Rep. 16, 897–906.

Deczkowska, A., Weiner, A., and Amit, I. (2020). The Physiology, Pathology, and Potential Therapeutic Applications of the TREM2 Signaling Pathway. Cell 181, 1207–1217.

Devitt, A., Moffatt, O.D., Raykundalia, C., Capra, J.D., Simmons, D.L., and Gregory, C.D. (1998). Human CD14 mediates recognition and phagocytosis of apoptotic cells. Nature 392, 505–509.

Edelstein, A.D., Tsuchida, M.A., Amodaj, N., Pinkard, H., Vale, R.D., and Stuurman, N. (2014). Advanced methods of microscope control using μManager software. J Biol Methods 1. https://doi.org/10.14440/jbm.2014.36.

Ellett, F., Pase, L., Hayman, J.W., Andrianopoulos, A., and Lieschke, G.J. (2011). mpeg1 promoter transgenes direct macrophage-lineage expression in zebrafish. Blood 117, e49–e56.

Elliott, M.R., Koster, K.M., and Murphy, P.S. (2017). Efferocytosis Signaling in the Regulation of Macrophage Inflammatory Responses. The Journal of Immunology 198, 1387–1394.

Fadok, V.A., Bratton, D.L., Konowal, A., Freed, P.W., Westcott, J.Y., and Henson, P.M. (1998). Macrophages that have ingested apoptotic cells in vitro inhibit proinflammatory cytokine production through autocrine/paracrine mechanisms involving TGF-beta, PGE2, and PAF. J. Clin. Invest. 101, 890–898.

Feng, M., Jiang, W., Kim, B.Y.S., Zhang, C.C., Fu, Y.-X., and Weissman, I.L. (2019). Phagocytosis checkpoints as new targets for cancer immunotherapy. Nat. Rev. Cancer 19, 568–586.

Feuerbach, D., Schindler, P., Barske, C., Joller, S., Beng-Louka, E., Worringer, K.A., Kommineni, S., Kaykas, A., Ho, D.J., Ye, C., et al. (2017). ADAM17 is the main sheddase for the generation of human triggering receptor expressed in myeloid cells (hTREM2) ectodomain and cleaves TREM2 after Histidine 157. Neuroscience Letters 660, 109–114. https://doi.org/10.1016/j.neulet.2017.09.034.

Freeman, S.A., and Grinstein, S. (2014). Phagocytosis: receptors, signal integration, and the cytoskeleton. Immunol. Rev. 262, 193–215.

Gheibi Hayat, S.M., Bianconi, V., Pirro, M., and Sahebkar, A. (2019). Efferocytosis: molecular mechanisms and pathophysiological perspectives. Immunol. Cell Biol. 97, 124–133.

Gumienny, T.L., Brugnera, E., Tosello-Trampont, A.C., Kinchen, J.M., Haney, L.B., Nishiwaki, K., Walk, S.F., Nemergut, M.E., Macara, I.G., Francis, R., et al. (2001). CED-12/ELMO, a novel member of the CrkII/Dock180/Rac pathway, is required for phagocytosis and cell migration. Cell 107, 27–41.

Hamieh, M., Dobrin, A., Cabriolu, A., van der Stegen, S.J.C., Giavridis, T., Mansilla-Soto, J., Eyquem, J., Zhao, Z., Whitlock, B.M., Miele, M.M., et al. (2019). CAR T cell trogocytosis and cooperative killing regulate tumour antigen escape. Nature 568, 112–116.

Hawley, K.L., Olson, C.M., Jr, Iglesias-Pedraz, J.M., Navasa, N., Cervantes, J.L., Caimano, M.J., Izadi, H., Ingalls, R.R., Pal, U., Salazar, J.C., et al. (2012). CD14 cooperates with complement receptor 3 to mediate MyD88-independent phagocytosis of Borrelia burgdorferi. Proc. Natl. Acad. Sci. U. S. A. 109, 1228–1232.

Heckmann, B.L., Boada-Romero, E., Cunha, L.D., Magne, J., and Green, D.R. (2017). LC3-Associated Phagocytosis and Inflammation. J. Mol. Biol. 429, 3561–3576.

Herbomel, P., Thisse, B., and Thisse, C. (2001). Zebrafish early macrophages colonize cephalic mesenchyme and developing brain, retina, and epidermis through a M-CSF receptor-dependent invasive process. Dev. Biol. 238, 274–288.

Hoijman, E., Häkkinen, H.-M., Tolosa-Ramon, Q., Jiménez-Delgado, S., Wyatt, C., Miret-Cuesta, M., Irimia, M., Callan-Jones, A., Wieser, S., and Ruprecht, V. (2021). Cooperative epithelial phagocytosis enables error correction in the early embryo. Nature 590, 618–623.

Hsiao, C.-C., van der Poel, M., van Ham, T.J., and Hamann, J. (2019). Macrophages Do Not Express the Phagocytic Receptor BAI1/ADGRB1. Front. Immunol. 0. https://doi.org/10.3389/fimmu.2019.00962.

Huang, J.F., Yang, Y., Sepulveda, H., Shi, W., Hwang, I., Peterson, P.A., Jackson, M.R., Sprent, J., and Cai, Z. (1999). TCR-Mediated internalization of peptide-MHC complexes acquired by T cells. Science 286, 952–954.

Janda, E., Boi, L., and Carta, A.R. (2018). Microglial Phagocytosis and Its Regulation: A Therapeutic Target in Parkinson’s Disease? Front. Mol. Neurosci. 11, 144.

Jaumouillé, V., Cartagena-Rivera, A.X., and Waterman, C.M. (2019). Coupling of β2 integrins to actin by a mechanosensitive molecular clutch drives complement receptor-mediated phagocytosis. Nat. Cell Biol. 21, 1357–1369.

Jay, T.R., von Saucken, V.E., and Landreth, G.E. (2017). TREM2 in Neurodegenerative Diseases. Mol. Neurodegener. 12, 56.

Kagan, J.C. (2017). Lipopolysaccharide Detection across the Kingdoms of Life. Trends Immunol. 38, 696–704.

Kamber, R.A., Nishiga, Y., Morton, B., Banuelos, A.M., Barkal, A.A., Vences-Catalán, F., Gu, M., Fernandez, D., Seoane, J.A., Yao, D., et al. (2021). Inter-cellular CRISPR screens reveal regulators of cancer cell phagocytosis. Nature https://doi.org/10.1038/s41586-021-03879-4.

Kaplan, G. (1977). Differences in the mode of phagocytosis with Fc and C3 receptors in macrophages. Scand. J. Immunol. 6, 797–807.

Kim, O.-H., Kang, G.-H., Hur, J., Lee, J., Jung, Y., Hong, I.-S., Lee, H., Seo, S.-Y., Lee, D.H., Lee, C.S., et al. (2022). Externalized phosphatidylinositides on apoptotic cells are eat-me signals recognized by CD14. Cell Death Differ. https://doi.org/10.1038/s41418-022-00931-2.

Kinoshita, T. (2020). Biosynthesis and biology of mammalian GPI-anchored proteins. Open Biol. 10, 190290.

Kojima, Y., Volkmer, J.-P., McKenna, K., Civelek, M., Lusis, A.J., Miller, C.L., Direnzo, D., Nanda, V., Ye, J., Connolly, A.J., et al. (2016). CD47-blocking antibodies restore phagocytosis and prevent atherosclerosis. Nature 536, 86–90.

Lam, W.A., Rosenbluth, M.J., and Fletcher, D.A. (2007). Chemotherapy exposure increases leukemia cell stiffness. Blood 109, 3505–3508.

Lamers, C., Plüss, C.J., and Ricklin, D. (2021). The Promiscuous Profile of Complement Receptor 3 in Ligand Binding, Immune Modulation, and Pathophysiology. Front. Immunol. 12, 662164.

Li, M., and Yu, Y. (2021). Innate immune receptor clustering and its role in immune regulation. J. Cell Sci. 134. https://doi.org/10.1242/jcs.249318.

Matlung, H.L., Babes, L., Zhao, X.W., van Houdt, M., Treffers, L.W., van Rees, D.J., Franke, K., Schornagel, K., Verkuijlen, P., Janssen, H., et al. (2018). Neutrophils Kill Antibody-Opsonized Cancer Cells by Trogoptosis. Cell Rep. 23, 3946–3959.e6.

van der Merwe, P.A., and Dushek, O. (2011). Mechanisms for T cell receptor triggering. Nat. Rev. Immunol. 11, 47–55.

Miyanishi, M., Tada, K., Koike, M., Uchiyama, Y., Kitamura, T., and Nagata, S. (2007). Identification of Tim4 as a phosphatidylserine receptor. Nature 450, 435–439.

Moffatt, O.D., Devitt, A., Bell, E.D., Simmons, D.L., and Gregory, C.D. (1999). Macrophage recognition of ICAM-3 on apoptotic leukocytes. J. Immunol. 162, 6800–6810.

Molgora, M., Esaulova, E., Vermi, W., Hou, J., Chen, Y., Luo, J., Brioschi, S., Bugatti, M., Omodei, A.S., Ricci, B., et al. (2020). TREM2 Modulation Remodels the Tumor Myeloid Landscape Enhancing Anti-PD-1 Immunotherapy. Cell 182, 886–900.e17.

Morgens, D.W., Deans, R.M., Li, A., and Bassik, M.C. (2016). Systematic comparison of CRISPR/Cas9 and RNAi screens for essential genes. Nat. Biotechnol. 34, 634–636.

Morgens, D.W., Wainberg, M., Boyle, E.A., Ursu, O., Araya, C.L., Tsui, C.K., Haney, M.S., Hess, G.T., Han, K., Jeng, E.E., et al. (2017). Genome-scale measurement of off-target activity using Cas9 toxicity in high-throughput screens. Nat. Commun. 8, 15178.

Padilla-Rodriguez, M., Parker, S.S., Adams, D.G., Westerling, T., Puleo, J.I., Watson, A.W., Hill, S.M., Noon, M., Gaudin, R., Aaron, J., et al. (2018). The actin cytoskeletal architecture of estrogen receptor positive breast cancer cells suppresses invasion. Nat. Commun. 9. https://doi.org/10.1038/s41467-018-05367-2.

Park, S.-Y., and Kim, I.-S. (2017). Engulfment signals and the phagocytic machinery for apoptotic cell clearance. Exp. Mol. Med. 49, e331.

Park, D., Tosello-Trampont, A.-C., Elliott, M.R., Lu, M., Haney, L.B., Ma, Z., Klibanov, A.L., Mandell, J.W., and Ravichandran, K.S. (2007). BAI1 is an engulfment receptor for apoptotic cells upstream of the ELMO/Dock180/Rac module. Nature 450, 430–434.

Penberthy, K.K., and Ravichandran, K.S. (2016). Apoptotic cell recognition receptors and scavenger receptors. Immunol. Rev. 269, 44–59.

Peri, F., and Nüsslein-Volhard, C. (2008). Live imaging of neuronal degradation by microglia reveals a role for v0-ATPase a1 in phagosomal fusion in vivo. Cell 133, 916–927.

Ralston, K.S., Solga, M.D., Mackey-Lawrence, N.M., Somlata, Bhattacharya, A., and Petri, W.A., Jr (2014). Trogocytosis by Entamoeba histolytica contributes to cell killing and tissue invasion. Nature 508, 526–530.

Reimand, J., Arak, T., Adler, P., Kolberg, L., Reisberg, S., Peterson, H., and Vilo, J. (2016). g:Profiler—a web server for functional interpretation of gene lists (2016 update). Nucleic Acids Res. 44, W83–W89.

Schlepckow, K., Monroe, K.M., Kleinberger, G., Cantuti-Castelvetri, L., Parhizkar, S., Xia, D., Willem, M., Werner, G., Pettkus, N., Brunner, B., et al. (2020). Enhancing protective microglial activities with a dual function TREM2 antibody to the stalk region. EMBO Mol. Med. 12, e11227.

Su, S., Zhao, J., Xing, Y., Zhang, X., Liu, J., Ouyang, Q., Chen, J., Su, F., Liu, Q., and Song, E. (2018). Immune Checkpoint Inhibition Overcomes ADCP-Induced Immunosuppression by Macrophages. Cell 175, 442–457.e23. https://doi.org/10.1016/j.cell.2018.09.007.

Takahashi, K., Rochford, C.D.P., and Neumann, H. (2005). Clearance of apoptotic neurons without inflammation by microglial triggering receptor expressed on myeloid cells-2. J. Exp. Med. 201, 647–657.

Thomas, L., Bielemeier, A., Lambert, P.A., Darveau, R.P., Marshall, L.J., and Devitt, A. (2013). The N-terminus of CD14 acts to bind apoptotic cells and confers rapid-tethering capabilities on non-myeloid cells. PLoS One 8, e70691.

Underhill, D.M., and Goodridge, H.S. (2012). Information processing during phagocytosis. Nat. Rev. Immunol. 12, 492–502.

Van der Meeren, L., Verduijn, J., Krysko, D.V., and Skirtach, A.G. (2020). AFM Analysis Enables Differentiation between Apoptosis, Necroptosis, and Ferroptosis in Murine Cancer Cells. iScience 23, 101816.

Vorselen, D., Labitigan, R.L.D., and Theriot, J.A. (2020a). A mechanical perspective on phagocytic cup formation. Curr. Opin. Cell Biol. 66, 112–122.

Vorselen, D., Wang, Y., de Jesus, M.M., Shah, P.K., Footer, M.J., Huse, M., Cai, W., and Theriot, J.A. (2020b). Microparticle traction force microscopy reveals subcellular force exertion patterns in immune cell-target interactions. Nat. Commun. 11, 20.

Vorselen, D., Barger, S.R., Wang, Y., Cai, W., Theriot, J.A., Gauthier, N.C., and Krendel, M. (2021). Phagocytic “teeth” and myosin-II “jaw” power target constriction during phagocytosis. Elife 10. https://doi.org/10.7554/eLife.68627.

Walbaum, S., Ambrosy, B., Schütz, P., Bachg, A.C., Horsthemke, M., Leusen, J.H.W., Mócsai, A., and Hanley, P.J. (2021). Complement receptor 3 mediates both sinking phagocytosis and phagocytic cup formation via distinct mechanisms. J. Biol. Chem. 296, 100256.

Wang, Y., Cella, M., Mallinson, K., Ulrich, J.D., Young, K.L., Robinette, M.L., Gilfillan, S., Krishnan, G.M., Sudhakar, S., Zinselmeyer, B.H., et al. (2015). TREM2 lipid sensing sustains the microglial response in an Alzheimer’s disease model. Cell 160, 1061–1071.

Wang, Y., Zhang, X., and Cai, W. (2019). Spherical harmonics method for computing the image stress due to a spherical void. J. Mech. Phys. Solids 126, 151–167.

Wright, S.D., Ramos, R.A., Tobias, P.S., Ulevitch, R.J., and Mathison, J.C. (1990). CD14, a receptor for complexes of lipopolysaccharide (LPS) and LPS binding protein. Science 249, 1431–1433.

Yeh, F.L., Hansen, D.V., and Sheng, M. (2017). TREM2, Microglia, and Neurodegenerative Diseases. Trends Mol. Med. 23, 512–533.

Yin, C., and Heit, B. (2021). Cellular Responses to the Efferocytosis of Apoptotic Cells. Frontiers in Immunology 12. https://doi.org/10.3389/fimmu.2021.631714.

Zhao, S., Zhang, L., Xiang, S., Hu, Y., Wu, Z., and Shen, J. (2022). Gnawing Between Cells and Cells in the Immune System: Friend or Foe? A Review of Trogocytosis. Front. Immunol. 13, 791006.

