## Supplement for "Cell surface receptors TREM2, CD14 and integrin α_M_β_2_ drive sinking engulfment in phosphatidylserine-mediated phagocytosis"

### Affiliations

### Supplemental Figure S1

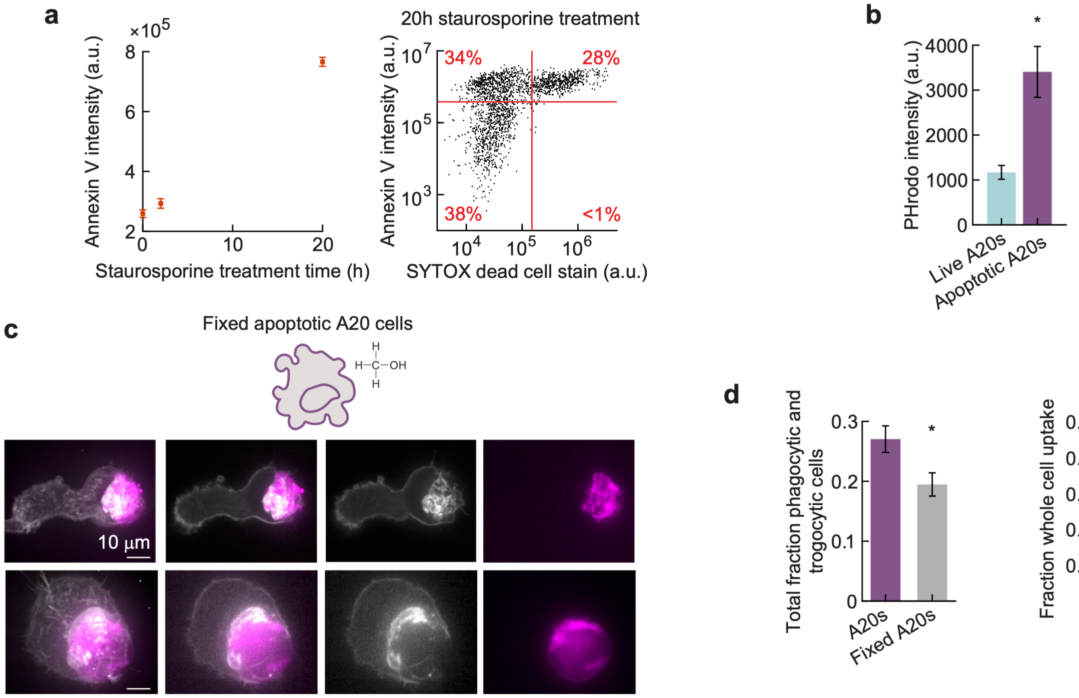


**Supplemental Figure S1: Phagocytosis of staurosporine-treated apoptotic cells. a,** Phosphatidylserine exposure upon staurosporine (STS) treatment (5 μM) of A20 cells, as evaluated using Annexin-V Alexa Fluor 647 and flow cytometry. Right: Labeling with both annexin V and the nuclear dead cell stain SYTOX orange allows distinguishing live cells (lower left, 38%), apoptotic cells (upper left; 34%) and dead cells (upper right; 28%). **b,** Uptake of live and apoptotic A20s (5 μM STS-treated for 20 h). A20s were stained using the pH-sensitive pHrodo-red dye that is quenched at neutral pH, and is brightly fluorescent in acidified phagosomes. Error bars indicate standard error of the mean of 3 independent experiments. *p = 0.04 as determined by a two-sample two-sided t-test. **c,** Uptake of apoptotic A20 cells that were treated with staurosporine similarly to **a**, and then briefly (15 s) fixed with methanol. Images acquired 30 minutes after bringing cells in close proximity. **d**, Left: quantification of total fraction of J774s with engulfed apoptotic cells or fractions thereof (left), evaluated after 90 minutes by evaluation of microscopy images. *p = 0.03 Right: Quantification of whole apoptotic cell uptake as a fraction of all partial and whole cell uptake events. ***p =$1.3\times10^{-7}$. All statistical tests were two-sided Wilcoxon rank-sum tests unless indicated otherwise.

### Supplemental Figure S2

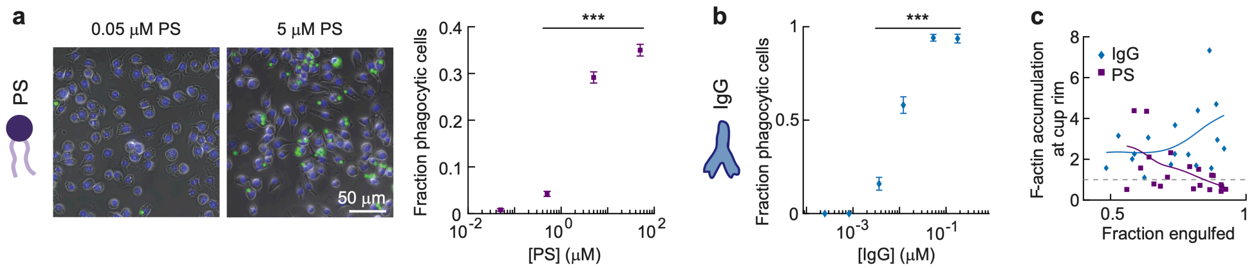


**Supplemental Figure S2. Ligand-dependence and F-actin accumulation during particle uptake. a,** Left: Composite epi-fluorescence and phase-contrast images (gray) of fixed J774 macrophages after 1 h incubation with 6.5 kPa DAAM-particles coated with various amounts of phosphatidylserine (PS) and TMR-cadaverine for visualization (shown in green). Cell nuclei were stained with Hoechst (blue). Right; quantification of fraction phagocytic cells for various PS-coating concentrations from microscopy images. At least 1,000 cells were analyzed in each condition. Markers show means and error bars indicate standard deviation (s.d.) estimated by treating phagocytosis as a Bernoulli process. Fractions were compared with the lowest PS concentration sample (0.05 μM) using Fisher's exact test (***$p < 10^{-7}$). **b,** Quantification of fraction phagocytic cells after 30 min incubation with 6.5 kPa DAAM-particles coated with various IgG concentrations. Line and marker styles as in **a**. At least 100 cells were analyzed for each condition (***$p < 10^{-6}$). **c,** Dependence of F-actin accumulation at the cup rim on phagocytic cup stage. Markers are individual phagocytic events, and lines are cubic smoothing splines indicating the trend of the data.

### Supplemental Figure S3

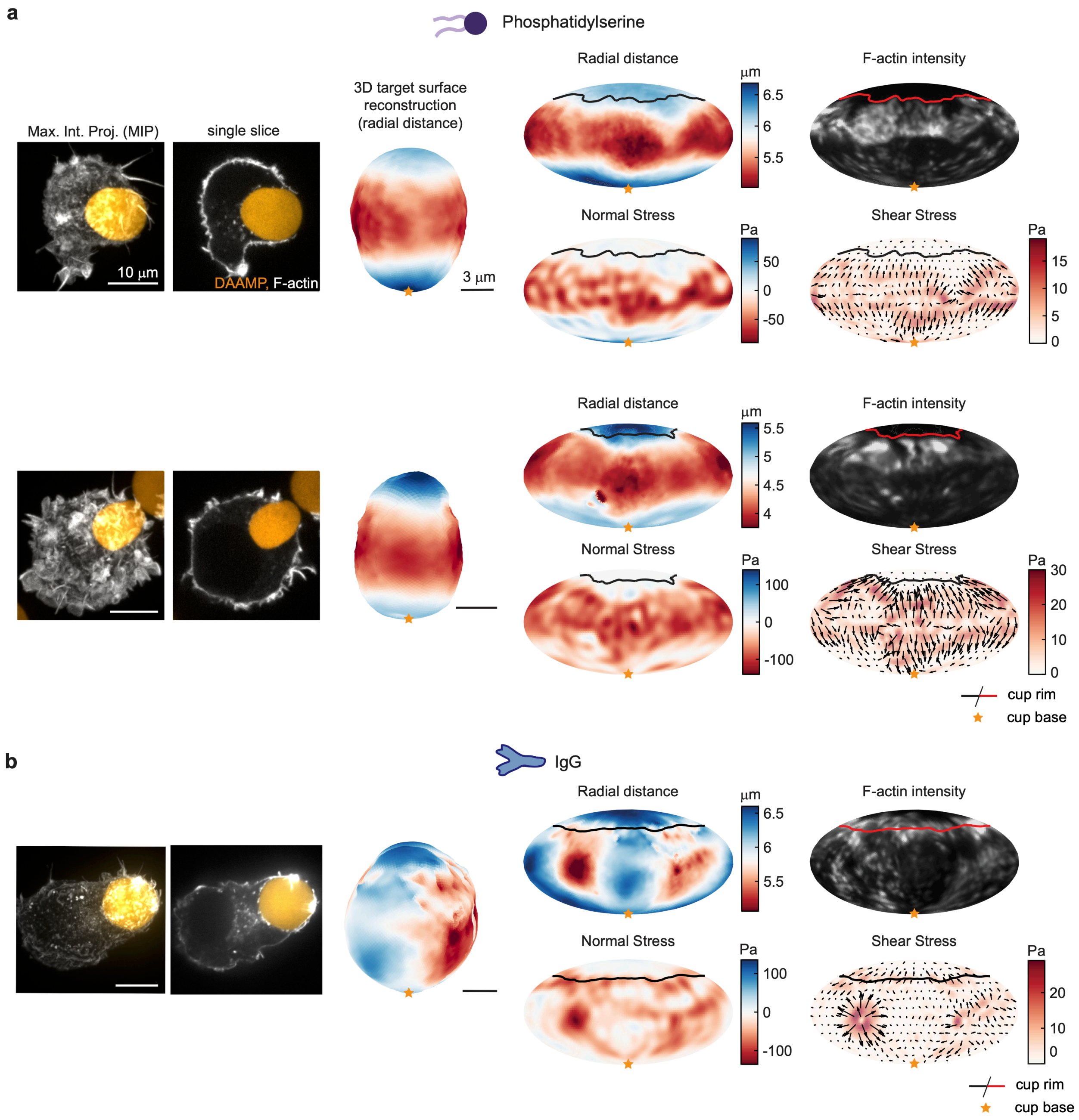


**Supplemental Figure S3: PS- and IgG-mediated phagocytosis of soft targets reveals cellular force exertion patterns. a,** Left: Confocal images of fixed J774 macrophages phagocytosing 0.3 kPa DAAM-particles functionalized with PS, and TMR-Cadaverine for visualization. Cells were stained for F-actin using phalloidin Alexa Fluor 488. Middle: 3D shape reconstructions of DAAM-particles revealing detailed target deformations induced during phagocytosis. Right: Mollweide projections showing target shape deformations, F-actin localization as well as normal and shear stresses inferred from target shape deformations over the entire target surface. Orange stars mark the base of the phagocytic cup, and cups are aligned with the phagocytic axis from the bottom to top. Force exertion in PS-mediated phagocytosis is mostly localized away from the cup rim and force exertion does not correlate strongly with F-actin localization. **b,** Similar to **a,** but for IgG-coated DAAM-particles of the same (0.3 kPa) rigidity. Although global target deformations are also observed in IgG-cups, F-actin and forces are distributed differently, with strong localization right at the rim of the cup. Moreover, the magnitude of forces correlates strongly with F-actin intensity and distinct protrusive spots, which have also been observed previously (Vorselen et al., 2020b, 2021)**,** can be observed within the cup rim.

### Supplemental Figure S4

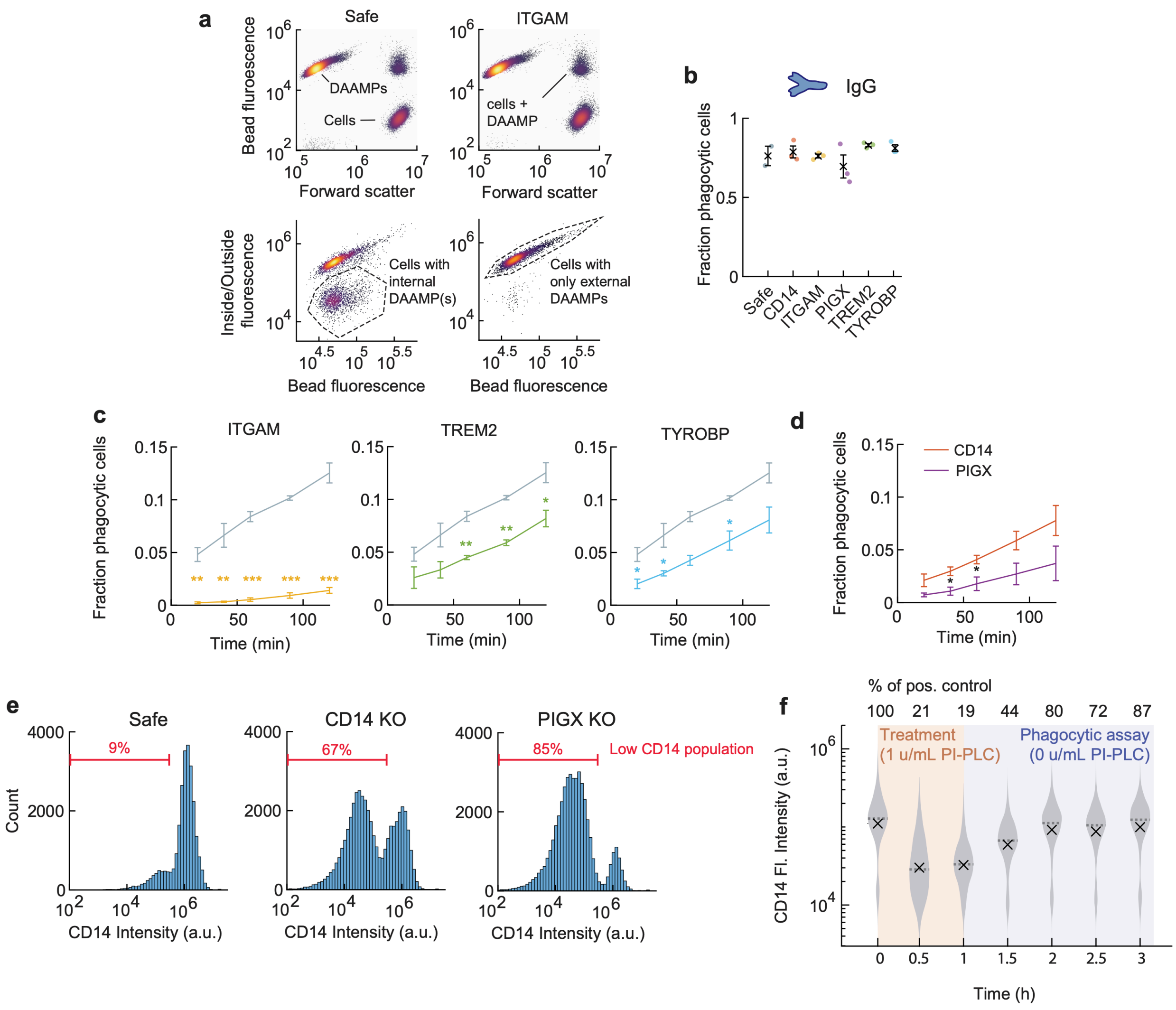


**Supplemental Figure S4: Quantification of phagocytosis by KO lines for PIGX, putative PS receptors, and PI-PLC treated cells. a,** Quantification of J774 macrophage phagocytic efficiency of DAAM-particles functionalized with PS, and TRITC-cadaverine for visualization, by flow cytometry. Experiments were carried out in suspension. Cells were fixed and the exposed DAAM-particle surface was immunostained. Top row: Density plots showing how DAAM-particles can be identified by their fluorescence, and cells by their higher forward scattering than DAAM-particles, which have very low refractive index because of their hydrogel nature. Bottom row: Gated data on the “cells + DAAM-particle(s)” population, where cells with an internal DAAM-particle can now be identified based on lack of signal of the exposed surface immunostaining (*i.e.* inside/outside stain). **b,** KO of PIGX and putative PS receptor doesn’t affect IgG-mediated phagocytosis. J774 macrophages were exposed to DAAM-particles functionalized with IgG for 60 minutes in solution. Individual markers indicate results of three replicate experiments, whereas black crosses and error bars indicate mean and s.d. of these experiments. No significant differences between the conditions were detected using two-sided two-sample t-tests, with at least 10,000 cells were analyzed for each cell line. **c,** Phagocytic efficiency of PS DAAM-particles for putative PS receptor KO lines determined in solution. Colored lines indicate the cell lines indicated above each graph. In total, at least $4\times10^5$ cells were analyzed for each cell line. Error bars indicate standard deviation (s.d.) of three replicate experiments. Two-sample two-sided t-tests were used to compare KO lines with safe guide KO control cells at each timepoint, with significance levels: $p < 0.05$*; $p < 0.01$**; $p < 0.001$***. Same data as represented in main text Fig. 5B. **d,** Direct comparison of uptake efficiency by CRISPR KO lines PIGX and CD14 measured by flow cytometry. This difference may be partially caused by differences in KO efficiency (see **e**). Error bars indicate standard deviation (s.d.) of three replicate experiments. In total, at least $4\times10^5$ cells were analyzed for each cell line. Data is also represented in main text Figs. 4B and 5B. Two-sample two-sided t-tests were used to compare KO lines at each timepoint, with significance level: $p < 0.05$*. **e,** CD14 and PIGX KO efficiency evaluated using CD14 immunostaining and evaluated by flow cytometry, indicating a higher KO efficiency of the PIGX line than the CD14 line. **f,** Efficiency of PI-PLC treatment evaluated by CD14 immunostaining, where shaded regions indicate CD14 exposure during treatment (0 - 1 h, orange) and after treatment has stopped (1 - 3 h, purple). Violin plots indicate the distribution of cells ($n > 3000$ for each timepoint), with mean (black cross) and median (dashed line).

### Supplemental Figure S5

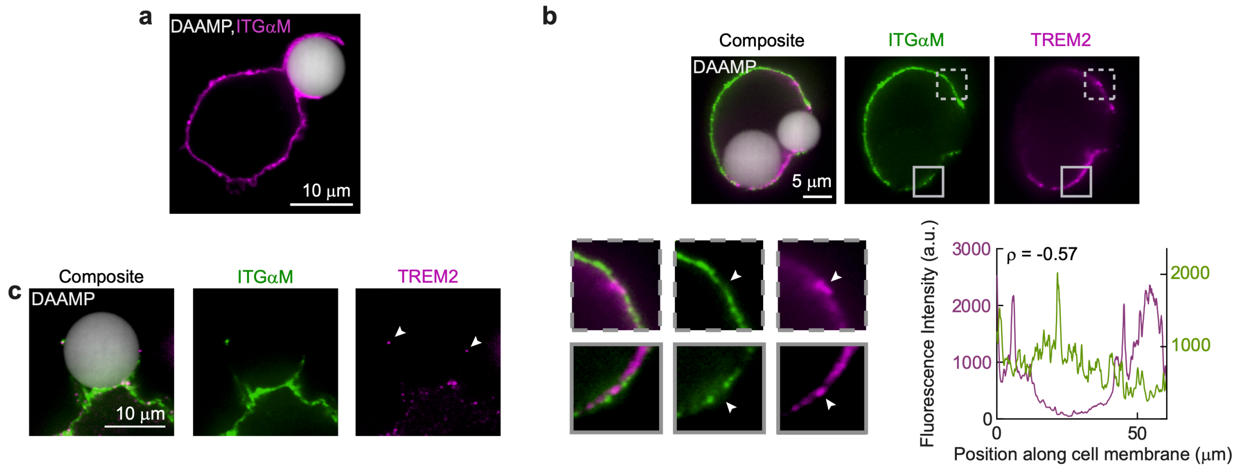


**Supplemental Figure S5: Immunofluorescent imaging of putative receptors during phagocytosis. a,** iSIM image of fixed J774 macrophages during engulfment of 6.5 kPa DAAM-particles functionalized with PS, and TMR-Cadaverine for visualization. Cells were immunostained for ITGαM. This rare outgrowing PS-cup shows high enrichment of ITGαM. **b,** iSIM image of fixed J774 macrophages during engulfment of 6.5 kPa DAAM-particles functionalized with PS, and TMR-Cadaverine for visualization. Cells were immunostained for ITGαM and TREM2, whose localization appears anti-correlated at the cell-scale (top row), but also more locally (zoomed images on the bottom left). Bottom right: Quantification of fluorescence along the cell border reveals a strong anti-correlation in localization. **c,** Similar to b, but showing TREM2 localization at the tips of “fingers” protruding around the target (highlighted with white arrowheads).

### Supplemental Table Titles and Legends

**Supplemental Table 1. Complete results for genome-wide PS-mediated phagocytosis screen in J774 macrophages.**

CasTLE analysis of genome-wide CRISPR knockout screen in J774 macrophages for phagocytosis of PS-coated DAAM particles. Includes CasTLE *P*-values, CasTLE scores, and CasTLE effect sizes for each gene, as well as hits defined using 1% FDR threshold. Two biologically independent screen replicates.

**Supplemental Table 2. Complete results for genome-wide IgG-mediated phagocytosis screen in J774 macrophages.**

CasTLE analysis of genome-wide CRISPR knockout screen in J774 macrophages for phagocytosis of IgG-coated DAAM particles. Includes CasTLE *P*-values, CasTLE scores, and CasTLE effect sizes for each gene, as well as hits defined using 1% FDR threshold. Two biologically independent screen replicates.
